# Phototropin modulates biosynthetic gene clusters network mediated biomanufacturing of anti-inflammatory metabolites in green lineage simply by illumination

**DOI:** 10.1101/2025.05.22.655495

**Authors:** Rajani Singh, Haseeb Ul Arfin, Amit Sharma, Nikita, Ravi Tandon, Peter Hegemann, Suneel Kateriya

**Affiliations:** Laboratory of Optobiotechnology, School of Biotechnology, Jawaharlal Nehru University, New Delhi 110067, India; Photo-Biology Laboratory, Multidisciplinary Centre for Advanced Research and Studies (MCARS), Jamia Millia Islamia (JMI), Jamia Nagar, New Delhi – 110025; Laboratory of AIDS Research and Immunology, School of Biotechnology, Jawaharlal Nehru University, New Delhi 110067, India; Institute of Biology, Experimental Biophysics, Humboldt-Universität zu Berlin, Berlin, Germany

**Author notes:** To whom correspondence should be addressed. Corresponding authors’. The authors contributed equally to this work (Rajani Singh, Haseeb Ul Arfin).

**Keywords:** Opto-biomanufacturing, biosynthetic gene clusters, carotenoid, photosynthesis machinery

## Abstract

Light sensing proteins, photoreceptor, coordinate with photosynthetic machinery, influencing both photosynthetic efficiency and several metabolic outcomes. This work deals with bottleneck issue of astaxanthin production from *Chlamydomonas reinhardtii.* We report a non-genetic, illumination-based strategy for biomanufacturing of astaxanthin under optimised blue light illumination in *C. reinhardtii.* Notably, first time ever, we identified and report astaxanthin synthesis-associated BGC in *C. reinhardtii* using plantiSMASH analysis. We discover experimental evidences for phototropin-modulated biosynthetic gene clusters (BGC)-mediated metabolite production in *C. reinhardtii*. We established that the prolonged exposure to fine-tuned blue light illumination substantially enhances the biomanufacturing of astaxanthin (1.6 times compared to red light) and pigments via the phototropin. It suggests that fine-tuned illumination conditions modulate molecular components in specific metabolite production. We are providing biochemical, genetic, transcriptomic, quantitative proteomics and systems biology evidences for opto-biomanufacturing of bioactive from green lineage via modulation of phototropin network with artificial illumination (without any genetic modification). By integrating data-driven analytics and systems biology-based computational pipeline, we elucidate the functional crosstalk between biosynthetic gene clusters with the phototropin and carotenoid metabolism synthesis pathway in the green lineage (terrestrial alga and higher plants). Thus, highlighting a new opto-biotechnology approach in metabolite production from various phototrophic organisms. These results establish a new avenue for opto-biomanufacturing strategies to generate bioactive molecules from microorganisms and other green lineages simply by illumination.

Highlights

- Opto-biomanufacturing of astaxanthin in *C. reinhardtii* is achieved simply by illumination.
- Biosynthetic gene clusters of metabolite production are also controlled by light.
- Photoreceptor-based opto-biomanufacturing of valuable bioactives is established.
- Opto-biomanufacturing opens avenues for bioactive production in various organisms.

## 1 Introduction

*Chlamydomonas reinhardtii* ability to grow at a large scale in a fermenter has recently been discovered, making it a suitable choice for the commercial production of high-value bioproducts. However, high biomass production and enhanced metabolite generation remain an unsolved challenge. Utilizing the photosynthetic efficiency of microalgae at its optimum might resolve these issues. Light, the most crucial parameter, affects algal physiology, altering algal biomass production and shaping their biochemical composition [1,2,3,4]. The coordinated light detection, utilization, and dissipation are vital for microalgal adaptation to environments with varying light intensity [5]. It spatiotemporally regulates the fitness of photosynthetic machinery [5]. Therefore, tailored illumination has potential for improved growth and enhanced algal metabolite accumulation. In *C. reinhardtii*, exposure to blue light of low intensity significantly upregulates the genes *phytoene desaturase*, *glutamate-1-semialdehyde aminotransferase*, and *light-harvesting polypeptides* (LHC), which are required for chlorophyll and carotenoid biosynthesis [6,7]. Moreover, the interactive effect of light and nutrients also affects photosynthesis and metabolite production [8]. Similarly, the light and temperature synergistic effect on growth kinetics and lipid accumulation was shown in *C. reinhardtii* [9]. Recently, a study provided detailed understanding of how blue light perception controls starch metabolism in *C. reinhardtii* [10]. Enhanced biomass production and biohydrogen accumulation were reported to be influenced by different light qualities and quantities [11]. Different studies have evidenced activation of downstream components of chlorophyll and carotenoid metabolism controlled by photoreceptor activity [6,7]. Hence, it is suggested that artificial modulation of algal photoreceptor(s) can alter biomass and change their biochemical makeup for improved manufacturing of valuable bioproducts. In *C. reinhardtii*, phototropin mediates regulation of photosynthesis [5], starch metabolism [10], influences expression levels of rhodopsins [12] and can modulate the expression of the genes involved in carotenoid synthesis [6]. Therefore, the photosynthetic (biomass) and metabolic (lutein, β-carotene, Chl a, Chl b, TAG, etc.) outputs from algae depend on the operational efficiencies of the photoreceptors involved in the algal photosynthetic and metabolic cycles. The enzymes and molecules pertaining to metabolic cycles are encoded in biosynthetic gene clusters. The biosynthetic gene clusters (BGCs) encodes interconnected genes, including operons and independently transcribed genes that collectively carry out the production of certain natural products [13]. However, photoreceptor-coordinated biosynthetic gene clusters (BGCs)-modulated metabolite production remains unexplored. Research towards carotenoid metabolism and metabolites related to this pathway mainly focused towards its overexpression via genetic engineering and high light stress. Moreover, so far, genetic engineering is mainly employed for astaxanthin (a valuable therapeutic metabolite) production in *C. reinhardtii* [14]. Researchers have shown that astaxanthin accumulation is widely accomplished by introducing β-carotene ketolase and high light stress [14, 15, 16]. As light governs the biomass and metabolites, here, we examined astaxanthin accumulation under different light regimes. Our studies prompted role of fined tuned illumination with photoreceptor coordinated networking for astaxanthin accumulation without genetic modification. Further, we uncover the underlying potential of light-regulated biosynthetic gene clusters for metabolite production in *C. reinhardtii*. Our proteomics data provided a proof of concept for light-regulated BGC-modulated metabolite production in green lineage without any genetic modification. This study establishes a sustainable, safe, and scalable paradigm for metabolite production without genetic manipulation, with strong implications for algal biotechnology, opto-biomanufacturing, and cascade biorefinery applications.

## 2 Methods

### 2.1 *In-silico* analysis for curated protein-protein networking between photosynthesis machinery, photoreceptors and different metabolic pathways

The crosstalk between enzymes of membrane lipid biosynthesis, photosynthetic pigments with photosynthesis machinery and photoreceptors in *C. reinhardtii* is shown by protein networking. The curated protein-protein network was predicted using String (https://string-db.org/) [17]. The results were visualised using Cytoscape 3.9.1 with default layout [18]. The network analysis was performed using the top 10 betweenness and shortest. Colour range (orange red to yellowish) indicates scores ranked by the betweenness method. Curated PPI networking was created based on literature, co-expression, and proteomics data and systems biology.

### 2.2 Bio-curation of opto-modulation of biosynthetic gene clusters (BGCs) for metabolite production

For bio-curation of opto-modulation of BGCs for metabolite production, BGCs were identified across different green lineages such as *C. reinhardtii* (freshwater algae), *Klebsormidium nitens* (terrestrial algae), *Arabidopsis thaliana* (higher plant, dicot) and *Oryza sativa* (higher plant, monocot). For identification of BGCs in *C. reinhardtii* and *K. nitens*, the genome sequence of each organism in fasta format and their annotation in GFF3 format were retrieved from NCBI. Further, this was uploaded in the plantiSMASH database and BGCs were identified using the default parameters and were compared with the already available BGCs in the plantiSMASH database. For BGCs annotation in *A. thaliana* and *O. sativa,* it is directly fetched from the plantiSMASH database [19]. For Bio-curation, curated protein networking showing crosstalk between BGCs, photoreceptor(s) and carotenoid metabolism were created using String database [17].

### 2.3 *Chlamydomonas* strain and culture conditions

The wild-type (CC-125) strain of *C. reinhardtii* and its phototropin CRISPR-Cas9 knockout mutant ΔPHOT-B5 mt+ were procured from the Chlamydomonas Resource Center (University of Minnesota) (https://www.chlamycollection.org/). The strain was maintained on TAP (Tris-Acetate-Phosphate) at 22 °C. Cells were initially grown on TAP (Tris-Acetate-Phosphate) agar plates (1.5% w/v). For experiments, cells were cultivated under a 12-hour light/12-hour dark photoperiod at a photon flux density (PFD) of 80 µmol m⁻² s⁻¹ using continuous warm white light (2700 K) in an incubator shaker maintained at 22°C. For liquid culture, a primary inoculum was prepared by resuspending cells in 50 mL of TAP medium supplemented with ampicillin (500 μg/ml) in a 250 mL Erlenmeyer flask. On the third or fourth day, when cultures reached the exponential growth phase, 1 mL of the primary culture was used to inoculate a 50 mL secondary culture in a 250 mL flask, without antibiotics. After three days, experimental cultures were established under the specified conditions (different light quality) for further analyses.

### 2.4 Growth curve analysis in *C. reinhardtii* under different illumination conditions

Experimental cultures were grown in a custom-designed multi-cultivator system comprising borosilicate glass tubes (120 mL capacity) containing 85 mL of culture medium. Cultures were exposed to monochromatic blue (450 nm), red (660 nm), and white light (2700 K) for a period of 3-4 days. Illumination was provided at a controlled photon flux density (PFD) of 120 µmol m⁻² s⁻¹. The photon flux density was measured using an Apogee Quantum Sensor (MQ-610). To ensure proper aeration and mixing, humidified 0.2 µm filtered air was bubbled uniformly into each tube. The temperature was maintained at 22 °C using a water bath system. The absorbance was taken at 680 nm and 750 nm every 24 hours using UV/Visible Spectrophotometer (BR Biochem. Life Science Pvt Ltd). Further, a growth curve was plotted as optical density vs. the number of days for each day. Moreover, the samples were pelleted at 4000 rpm for 10 minutes at 4 ℃ in the mid to late exponential phase for HPLC, qRT-PCR and proteomics analysis.

### 2.5 Cell morphology under different illumination conditions

*Chlamydomonas reinhardtii* cells (1 mL) were harvested by centrifugation in 1.5 mL microcentrifuge tubes, and the supernatant was discarded. The cell pellet was washed twice with 1X phosphate-buffered saline (PBS). Fixation was performed by resuspending the cells in 100 µL of 2.5% paraformaldehyde (PFA) (Sigma) and incubating them on ice in the dark for 20 minutes. The cells were then transferred to room temperature for 40 minutes while maintaining them in 2.5% PFA. Following fixation, the PFA was removed, and cells were washed thrice with PBS followed by resuspension in a final volume of 50 µL PBS.

For microscopy, 2 µL of the fixed cell suspension was placed on a clean coverslip, and a glass slide was gently inverted onto it, applying slight pressure. The coverslip was sealed using nail polish along the edges. Cells were visualized using a fluorescence microscope (Zeiss Axio Observer 7 with a 63x magnification oil immersion plan-apochromatic lens with NA 1.4) at 63× magnification with oil immersion. Imaging was conducted under the GFP channel with 15% intensity and an exposure time of 40 milliseconds, as well as under bright-field illumination with 8% intensity and a 20-millisecond exposure time.

### 2.6 Extraction and quantification of chlorophyll and carotenoid pigments in *C. reinhardtii* exposed to different light regimes

Extraction and quantification were performed by using both a UV-Vis spectrophotometer and HPLC analysis. Chlorophyll a, b and carotenoid contents were estimated using the Arnon et al. [20] method. Briefly, pellets were homogenized in 80 % 1 ml of acetone and centrifuged at 10000 rpm for 10 minutes. The optical density was recorded at 663 nm, 645 nm and 470 nm against 80 % acetone (blank) using Carry 3500 UV-Vis Compact Peltier, (Agilent; Australia). The concentrations of chlorophyll a, b, total chlorophyll and carotenoid were calculated using the Porro et al. [21] equation as described below

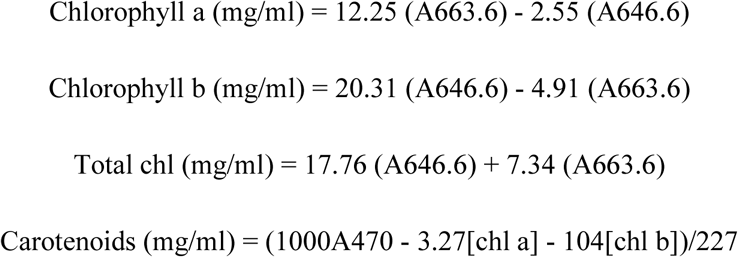

For mg/g, mg/mL amount is divided by the mass of the cells and multiplied by the total volume of the reaction.

For quantification of carotenoid pigments (lutein, astaxanthin, and zeaxanthin), HPLC was performed. Reversed-phase HPLC method was used for the quantification using ACQUITY BEHC 18 column with ACQUITY UPLC system (Waters Corp; USA). The extracts were partitioned into diethyl ether with a flow rate of 1 mL/minute, and the absorbance was recorded at 450 nm.

The astaxanthin content was determined by following method adapted from Sun et al [22]. The cells pellet were resuspended in methanol solution and heated for 5 min at 60 ℃ followed by centrifugation at 4000 rpm for 10 min. Next to the pellet, 2 mL DMSO was added and mixed properly to extract astaxanthin by centrifugation at 4000 rpm for 10 min. This step was repeated until the pellet turns white. The absorbance was measured after mixing all the supernatant at 475 nm using Carry 3500 UV-Vis Compact Peltier, (Agilent; Australia). The astaxanthin was calculated by absorbance coefficient 0.191 cm2/μg using formula

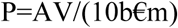

Where A is absorbance at 475 nm, V is total volume (mL), b, is the path length, m is the weight of the sample (mg) and € is the absorbance coefficient.

### 2.7 Bioassay to test the bio-compatibility and anti-inflammatory activity responses of the algal extract in macrophage RAW cell line

To check the potency of anti-inflammatory product response, first of all, MTT was performed to check the concentration of astaxanthin (which has anti-inflammatory activity) and algal extract on Raw 264.7 cells. The cytotoxic effects of astaxanthin (positive control) and the experimental extract were evaluated in RAW 264.7 murine macrophage cells using the MTT assay. RAW 264.7 cells were cultured in Dulbecco’s Modified Eagle Medium (DMEM) supplemented with 1% of antibiotic-antimycotic solution and 10% of fetal bovine serum (FBS) at standard culture conditions (37°C, 5% CO₂).

For the assay, cells were seeded in a 96-well culture plate at an appropriate density and allowed to adhere overnight. Astaxanthin and the extract were dissolved in acetone and further diluted in complete culture medium to obtain the required treatment concentrations. Corresponding acetone-treated wells were maintained as solvent controls. After incubation, the cells were treated with different concentrations of astaxanthin and the extract, and with an acetone control for 24 hours. Untreated cells served as the negative control. Following treatment, the medium was removed, and 5 mg/mL MTT reagent (prepared in PBS) was added to each well and kept at 37°C for 3–4 hours. Subsequently, the supernatant was carefully discarded, and the formed formazan crystals (obtained after incubation) were dissolved using DMSO (or appropriate solubilization solution). The absorbance was examined at 570 nm with the help of microplate reader.

Cell viability was calculated with respect to untreated control cells. All experiments were performed in triplicates, and the value represented as mean ± standard deviation. Statistical signification was checked by comparison between different treatment groups and control groups using GraphPad Prism software. One-way ANOVA followed by post hoc multiple comparison test was used to determine statistical significance among groups. Further, qRT- PCR was performed to assess the potency of anti-inflammatory product obtained. Actin was used as reference gene to normalize the gene expression. Melt curve analysis was also performed to ensure specificity of the expression.

### 2.8 Comparative quantification of different photoreceptor genes in *C. reinhardtii* under different light conditions using real-time quantitative PCR

For quantification of different photoreceptor gene expression, total RNA was extracted from wild-type algal culture grown till exponential phase (O.D. 0.5-0.7 at 750 nm) using the TRIzol method [23]. It is followed by cDNA synthesis using AffinityScript cDNA synthesis kit (Agilent, Santa Clara, USA) as per the manufacturer’s instructions. Independent qPCR reaction was carried out using Kappa^TM^ SYBR^R^ Fast mix as per the manufacturer’s protocol with Stratagene Mx3000P machine (Stratagene, La Jolla, CA, USA). The thermal profile employed was: 10 minutes at 95 °C, 40 cycles of denaturation for 10 seconds at 95 °C, annealing for 1 minute at 60 °C and extension for 1 minute at 72 °C. The CBLP housekeeping gene was used as the internal reference for normalization of DNA quality and quantity. The relative expression of photoreceptor(s) gene was determined by the ΔΔCt approach using the threshold value (Ct). At the end of the cycles, melt curve analysis was performed, ensuring a single specific product for each gene. The qPCR experiment was conducted using two biological and two technical replicates for each sample. The primers adapted from Causo et al. [24]. Further, for quantification of different photoreceptor proteins, immunoblot analysis performed following lab-established protocol [25].

### 2.9 Label-free quantitative proteomics under red and blue light in the whole cell proteome of *C. reinhardtii*

#### 2.9.1 Sample preparation for proteomics

The sample preparation and mass spectrometric analysis were performed by Vproteomics. At first, the protein sample (25 µg) was reduced by 5 mM TCEP, followed by alkylation using 50 mM iodoacetamide. Subsequently, the protein was digested with trypsin (trypsin-to-lysate ratio 1:50) at 37 °C for 16 hours. This is followed by purification using a C18 silica cartridge. The obtained mixture was dried in a speed vacuum to concentrate it. Further, resuspension of the dried pellet was done in buffer A containing acetonitrile (2%) and formic acid (0.1%)

#### 2.9.2 Mass spectrometric analysis of peptide mixtures

For mass spectrometric analysis, all the experiments were run on an Easy-nLC-1000 system (Thermo Fisher Scientific) coupled with an Orbitrap Exploris 240 mass spectrometer (Thermo Fisher Scientific). It is equipped with a nano-electrospray ion source. At first, about one µg of phosphopeptides sample was dissolved in buffer A (2% acetonitrile and 0.1% formic acid). It is then resolved using a Picofrit column (1.8-micron resin, 15 cm length). This was followed by gradient elution using a 0–38% gradient buffer B containing formic acid (0.1%) and acetonitrile (80%) at 500 nL/minute flow rate for 96 minutes, followed by 90% of buffer B for 11 minutes. At last, column equilibration was performed for 3 minutes. MS data acquisition was performed using Orbitrap Exploris 240. The conditions used were: Max IT = 60ms; AGC target = 300%; RF Lens = 70%; R = 60K, mass range = 375−1500. MS2 data was collected using the following conditions: Max IT= 60ms, R= 15K, AGC target 100%. MS/MS data acquisition was done using a dynamic data-dependent top20 method, which chooses the most abundant precursor ions from the survey scan, employing dynamic exclusion for 30 seconds.

#### 2.9.3 Data processing and Gene Ontology (GO) enrichment analysis

Vproteomics (New Delhi, India) performed the differential expressed protein identification. For the sample processing and RAW file generation, Proteome Discoverer (v2.5) against the Uniprot reference database was used. For dual Sequest and Amanda search, the precursor was set at 10 ppm, whereas fragment mass tolerances were set to 0.02 Da. Further, the protease was used to generate peptides. The carbamidomethyl on cysteine is considered a fixed modification, and oxidation of methionine and N-terminal acetylation are considered variable modifications for database search. The protein false discovery rate and peptide spectrum match were set to 0.01 FDR. The GO enrichment pathway analysis was done using Omicsbox tool (BioBam bioinformatics, S.L.) [26] and g:profiler tool [27].

### 2.10 Statistical analysis for data validation

Three independent biological replicates were used for each experiment. One-way ANOVA with Tukey’s test, and t-test were applied for statistical analysis of data using GraphPad Prism (version 5) software. The mean values significance level at P < 0.05, P < 0.001, P < 0.01 and P < 0.05 were denoted by ***, ** and * respectively, ns denotes non-significant changes.

## 3 Results

### 3.1 Biosynthetic gene cluster (BGCs) molecular components reveals photoreceptor-coordinated mechanism for BGCs-mediated carotenoid metabolism in *C. reinhardtii*

Photosynthesis in algae depends on the efficient functioning of several light-dependent components, including photoreceptors. Two or more photoreceptors act in a coordinated manner and exert a coalescent effect on metabolic pathways and photosynthetic efficiency [5]. Photoreceptor also affects the downstream components of carotenoid metabolism [6,7]. As molecular components of these biosynthetic pathways are clustered together as BGCs, we studied biosynthetic gene clusters in light of opto-biomanufacturing across different green lineage, a new avenue in opto-biotechnology. We therefore identified the BGCs present in *C. reinhardtii* (fresh water algae) and *Klebsormidium nitens* (terrestrial algae, an evolutionary link between lower and higher plants) using the plantiSMASH tool. A total of 3 BGC clusters were identified in *C. reinhardtii*, while 7 clusters were identified in *K. nitens* (Table 1, Additional data 1). Among them, most of the clusters encode the genes for fatty-acid-saccharide and fatty-acid-saccharide-alkaloid production in *C. reinhardtii*. Interestingly, we found a cluster encoding genes for β-carotene hydroxylase and β-carotene ketolase along with a gene encoding starch synthesis and a putative protein. In *K. nitens*, most of the cluster encodes for saccharide and another one is putative or polyketides (Table 1).

**Table 1:** List of different biosynthetic gene clusters identified in *C. reinhardtii* and *Klebsormidium nitens using* plantiSMASH.

| Clusters | Type | Size (kb) | Core domains |
| --- | --- | --- | --- |
| *Chlamydomonas reinhardtii* |  |  |  |
| Cluster1 | Fatty acid-saccharide-alkaloid | 27.11 | FA_desaturase,<br>Glycos_transf_1,p450 |
| Cluster2 | Putative | 58.96 | Transferase, p450 |
| Cluster3 | Fatty acid-saccharide | 49.59 | FA_desaturase, Glycos_transf,<br>FA_hydroxylase |
| *Klebsormidium nitens* |  |  |  |
| Cluster 7 | Saccharide | 68.11 | AMP-binding, Epimerase,<br>UDPGT |
| Cluster 6 | Polyketide | 80.25 | Chal_sti_synt_C,<br>NAD_binding_1, adh_short |
| Cluster 5 | Lignan-Saccharide | 61.42 | Dirigent, Epimerase,<br>Glycos_transf_2 |
| Cluster 4 | Saccharide | 81.61 | Amino_oxidase,<br>Glycos_transf_1, adh_short |
| Cluster 3 | Saccharide | 147.14 | Epimerase, FA_desaturase,<br>Glycos_transf_1 |
| Cluster 2 | Lignan | 54.59 | Dirigent, Epimerase,<br>Methyltransf_7 |
| Cluster 1 | Saccharide | 32.60 | Transferase, UDPGT, p450 |

As we are hypothesising that light might govern the expression of BGCs via photoreceptors, the bio-curated protein-protein networking for each identified cluster with photoreceptor(s) was created. The PPI shows crosstalk between proteins from the BGCs cluster and photoreceptor(s) (Figure 1A). Further, we have also performed an in-depth analysis for BGCs encoding fatty acid-saccharides and alkaloid. PPI networking implicated that the genes/proteins of two clusters showed indirect interactions with the different photoreceptors in *C. reinhardtii* (Figure 1B) while direct interaction of photoreceptor in *K. nitens* with BGCs was observed as shown in supplementary figure (Figure S1). We found that photoreceptor(s) phototropin, and rhodopsin showed direct interaction with glycosyl transferase family proteins (BGC core domain component) identified in *K. nitens.* Thus, this unveils a link between light and BGCs-mediated metabolite production.

**Figure 1:**
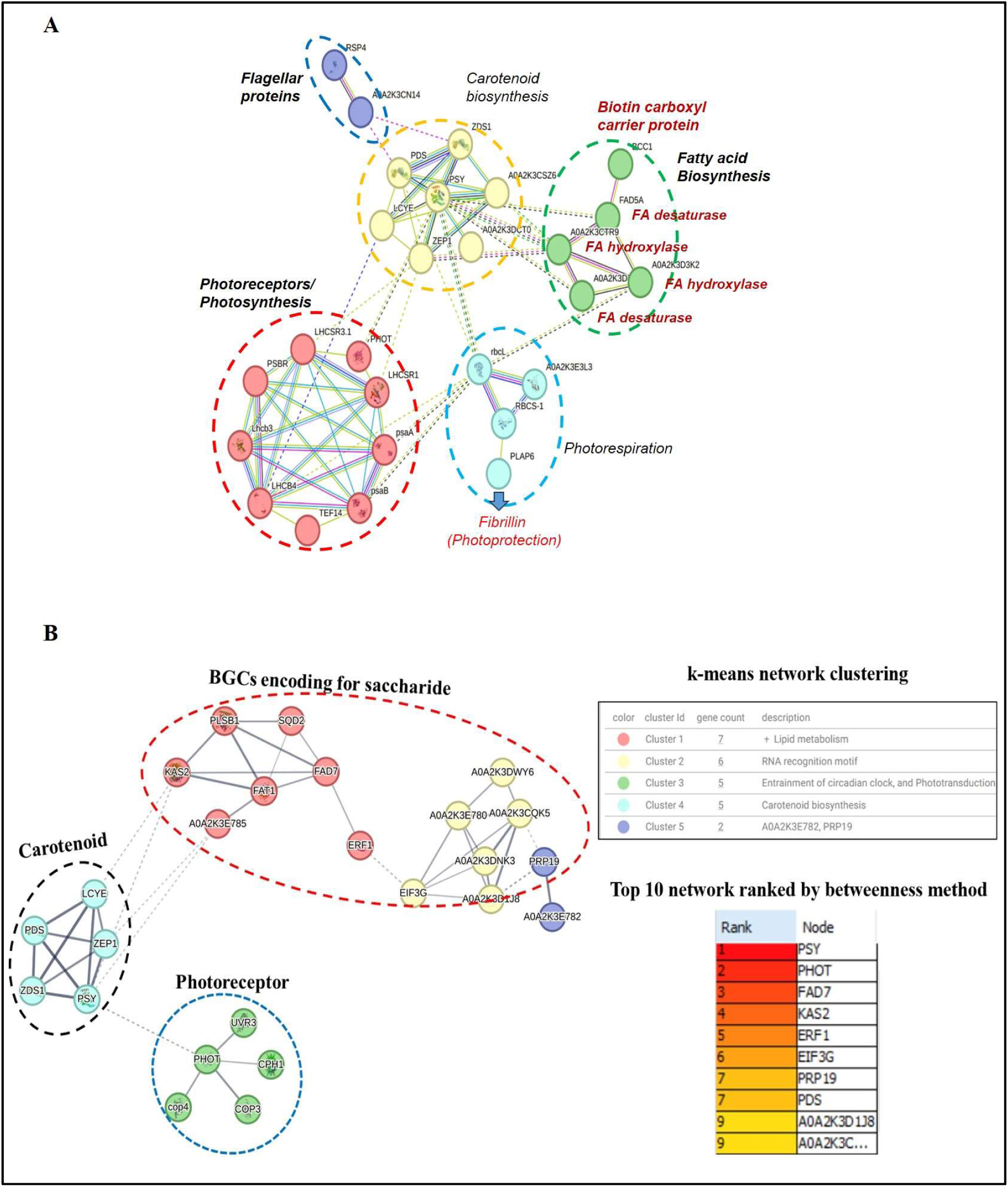
Bio-curation of opto-modulation of Biosynthetic Gene Cluster for targeted metabolite production. A: Represents the crosstalk of the molecular components of BGCs encoding fatty acid-saccharides-alkaloids with carotenoid metabolism in *C. reinhardtii*. Here, the FA desaturase and glycosyl-transferase show secondary interaction with photoreceptor (marked in a blue colour circle). Further, it also showed interaction with FA metabolism enzyme (marked in green circle). The CHYB shows interaction with PSY with is directly regulated by phototropin. The bio-curation of PPI networking was done using String database based on text mining, co-expression, experimental data and lab proteomics data. The list of important nodes regulating the network is given in supplementary file. **B: Shows the important nodes controlling the PPI network.** The k-means clustering method was applied to cluster the PPI into 5 clusters. All nodes that control the overall network were analysed using CytoHubba analysis using the betweenness algorithm and shortest path method. The scores ranked for interacting partners using the betweenness method are indicated in colour, ranging from red to yellow. SQD2- Uncharacterized protein, contains domain glycosyl_transferase_1-4; FAD7- Chloroplast glycerolipid omega-3-fatty acid desaturase; PHOT- Phototropin, Blue light-sensing protein; KAS2-3-oxoacyl-[acyl-carrier-protein] synthase; ERF1- Eukaryotic release factor 1; PDS-Phytoene desaturase; PRP19- Spliceosome component, nuclear pre-mRNA splicing factor; A0A2K3D1J8- Uncharacterized protein; EIF3G- Eukaryotic translation initiation factor 3 subunit G; RNA-binding component of the eukaryotic translation initiation factor 3 (eIF-3) complex, which is involved in protein synthesis of a specialized repertoire of mRNAs and, together with other initiation factors, stimulates binding of mRNA and methionyl-tRNAi to the 40S ribosome. The eIF-3 complex specifically targets and initiates translation of a subset of mRNAs involved in cell proliferation.

### 3.2 Photoreceptor(s) expression get influenced by prolonged blue and red light in *C. reinhardtii* at selected light regime

Blue light sensor phototropin is involved in maintaining the overall fitness of photosynthesis machinery and photo protection [5]. Earlier studied demonstrated that it regulates gene expression of proteins important in photosynthesis, photoprotection, gametogenesis, carotenoid metabolism and adhesiveness in *Chlamydomonas* [5, 6]. Phototropin expression is also influenced by red light [6]. We intended to study that tailored illumination condition’s effects on photoreceptor mediated metabolite production. To test the hypothesis, at first, we exposed dark-adapted synchronized cells to different light wavelength for 2 hours at approximately 4000 lux. The effect of different light conditions was examined on the basis of chlorophyll amount, carotenoid content and phototropin expression. The lights of different wavelengths (blue, red, green, white, UV and dark) at 4000 lux for 2 hours exposure showed varied effects on phototropin level (Figure 2A and B) and photosynthetic pigments in *C. reinhardtii* (see in supporting information Figure S2). Significant variation was observed in phototropin (phot) expression (1.5 and 1.2 times). The photosynthetic pigments (2-fold change and 3-fold change) in particularly red and blue light respectively, as compared to white when exposed for 2 hours at the end of the dark phase (Figure S2). Most of light-driven studies were based on exposure of light for few seconds to hours. Here we grown the culture under blue and red light (12 hours light/12 hours dark) till exponential phase.

**Figure 2:**
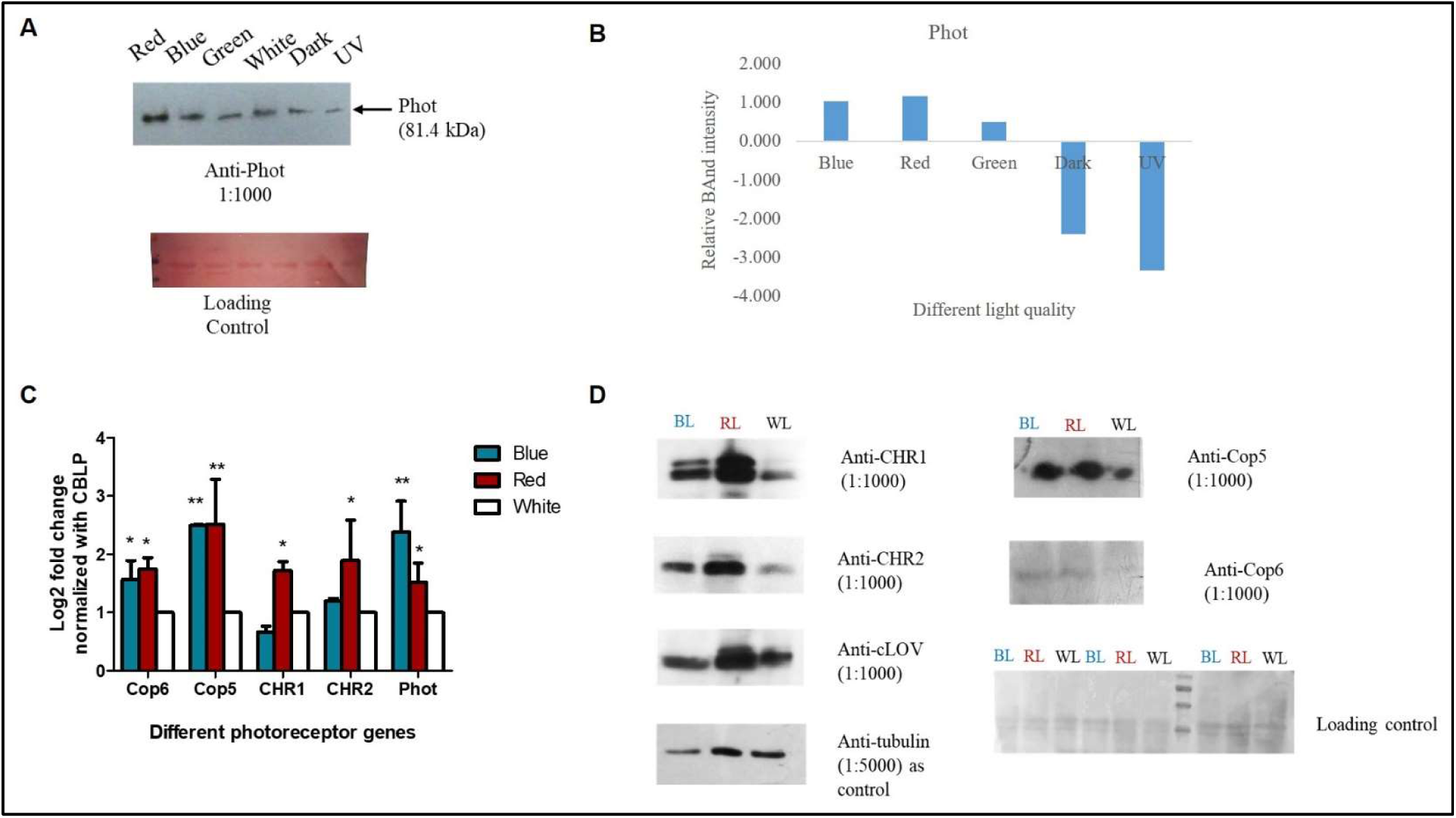
Represents the (A, B) Effect of different wavelengths of light on phototropin in *C. reinhardtii*. Immunoblot from CrTCL (*C. reinhardtii* total cell lysate) with anti-Phot (cLOV) with dilution 1:1000) antibody. © qRT-PCR analysis of different genes like *CHR1, CHR2, Phot, Cop5* and *Cop6* in *C. reinhardtii*, after blue light and red light treatment (Control samples-sample exposed in white light). Values are mean ± SE (n = 4). *P < 0.05, **P < 0.01, ***P < 0.001. (D) Immunoblot from CrTCL (*Chlamydomonas reinhardtii* total cell lysate) with anti-Phot (cLOV), anti-CHR1, CHR2, Cop5 and Cop6 antibodies with dilution 1:1000. Tubulin was used as control for Phot, CHR1 and CHR2 while ponceau was used as control for Cop5 and 6. BL- Blue light, RL- Red light, WL- white light

We next examined photoreceptor expression on prolonged diurnal blue and red light exposure. Here, in the present study, the expression of photoreceptors are also influenced by different light conditions (Figure 2C and D). A substantial increase in *phot, Cop5 and Cop6* gene expression was observed when exposed to red and blue light with respect to white. *ChR1* and *ChR2* expression were significantly upregulated under red light, whereas no change in their expression was observed under blue light illumination (Figure 2C and D). The red light-sensing protein, phytochrome, is not present in *Chlamydomonas*. Therefore, upregulation of phototropin in red light is due to the involvement of phototropin interaction with some other phytochrome-like proteins, putatively red light-sensitive protein (animal-like cryptochromes, aCRY) [28]. Recently, our group demonstrated phototropin interaction with other photoreceptors such as Chlamyopsin 3 (ChR1) and Chlamyopsin 4 (ChR2), Chlamyopsin 5 and 6 (Cop5 and Cop6) in C. *reinhardtii* [25]. Our result suggests a crucial role of illumination conditions on photoreceptor, in turn, photo-physiological responses in algae.

### 3.3 Upregulation of targeted metabolite content under the influence of prolonged blue light in *C. reinhardtii* without altering total protein level

Various light-driven studies were based on exposure of light for few seconds to hours. Here we grown the culture under blue and red light (12 hours light/12 hours dark) till exponential phase. So, we next examined the biochemical compositions of *C. reinhardtii* on the selected light regimes. Protein, chlorophyll and carotenoid content were assessed under different light regimes. No effect on protein amount was observed among different light spectra (Figure S2C). We reasoned that the protein level is also affected by the different growth phase, cell density along with light conditions. As the specific growth rate remains same in our conditions, protein amount remains same. Our finding corroborates with Yuan et al. [10]. Further, a significant increase in chlorophyll and carotenoid content was observed in blue and red light. Carotenoid content was substantially enhanced in the presence of blue light (Figure S2D, Additional data 3). A substantial 2-fold increase in Chl content was noted as compared to white and red light. Similarly, exposure to red light resulted in an apparent change (one-fold increase) in carotenoid content w.r.t. to white. However, the specific growth rates of *C. reinhardtii* cultures may appear similar under red, white, and blue light conditions, notable differences emerge during the stationary phase (Figure S3). Results indicated that different wavelengths (red, blue and white) of light significantly affected the algal physiology in terms of chlorophyll and carotenoid content. Thus, this suggests that light imparts a striking effect on metabolite production [3,29].

Besides this, continuous exposure to blue light (12 hours light/12 hours dark) for 3-4 days leads to enhanced accumulation of targeted carotenoid metabolites with respect to red light (Figure 3A, Additional data 4). A total of 1.3-fold increase in lutein and zeaxanthin amount was observed on blue light exposure. Several studies aligns with our results. Light quality regulates the lutein content in different algal species such as *Dunaliella salina* [30], *Chlorella vulgaris* [31], *Haematococcus pluvialis, Chlamydomonas sp JSC4* [32]. Moreover, notably, we also observed the accumulation of astaxanthin in *C. reinhardtii* simply by playing with the illumination condition.

**Figure 3:**
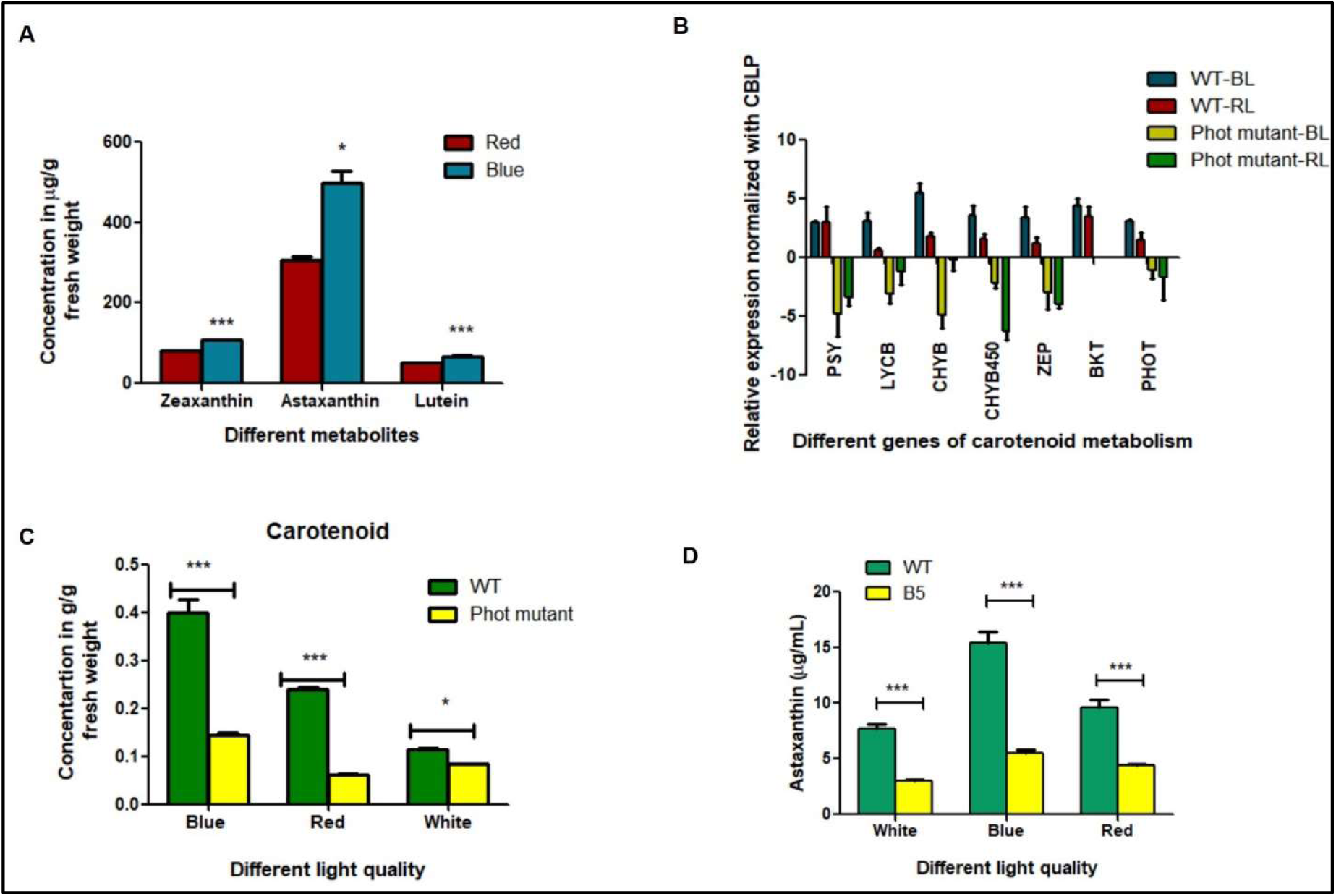
Effect of phototropin-mediated opto-modulation on metabolite gene expression and targeted metabolites (astaxanthin, zeaxanthin, and lutein) content in *C. reinhardtii* and its phototropin mutant (CRISPR-Cas9 knockout ΔPHOT-B5 mt+). (A): Effect of different light conditions on targeted metabolite content in *C. reinhardtii*. The zeaxanthin, astaxanthin, and lutein were quantified using HPLC at mid-exponential phase. The standard curve for each pigment is shown in Figure S4. (B): Represents the gene expression profile of *ZEP, PSY, PDS, LYCB, CHYB, BKT* and *bp450* in *C. reinhardtii* CRISPR-Cas9 phot mutant B5, after blue light and red-light treatment (Control samples-sample exposed to blue light). (C) Effect of different wavelengths of light on phototropin mutant carotenoid contents compared with its wild-type CC-125. (D) Represents the astaxanthin in *C. reinhardtii* CRISPR-Cas9 phot mutant B5, after blue light and red-light treatment. Values are mean ± SE (n = 4). *P < 0.05, **P < 0.01, ***P < 0.001, ns – non-significant changes. The t-test was performed for significance analysis. (Values are mean ± SE (n = 2, technical replicates and 2 biological replicates). PSY; Phytoene synthase, CHYB; beta-carotene hydroxylase, LYCB; lycopene β-cyclase, ZEP; zeaxanthin epoxidase, PDS; phytoene desaturase, BKT; beta-carotene ketolase and Bp450; P450 cytochrome–dependent β-ring carotene hydroxylase genes.

### 3.4 Astaxanthin is accumulated in *C. reinhardtii* simply by prolonged blue light illumination without genetic modification

Astaxanthin is an important and valuable metabolite. Due to its antioxidant and anti-inflammatory properties, it is widely used as an active ingredient in dietary supplements. Its enhanced production is mainly reported under high light stress in different microalgae [14,15]. However, studies demonstrated that *C. reinhardtii*, in native form does not make astaxanthin. Efforts have been made to accumulate astaxanthin in *C. reinhardtii*. *C. reinhardtii* is mainly genetically engineered for enhanced production of astaxanthin [14]. It is reported that astaxanthin only accumulates after genetically incorporating β-carotene ketolase (BKT) alone and in combination with β-carotene hydroxylase in *C. reinhardtii* [14,15,33]. Interestingly, here, for the first time, we obtained accumulation of astaxanthin in *C. reinhardtii* without any genetic modification simply by modulating light conditions (Figure 3A, S4). The chromatogram is given in Figure S4. Continuous blue and red-light illumination (12-hour light/12 hour dark photoperiodic at 120 µmol m⁻² s⁻¹) for 3-4 days resulted in the natural accumulation of astaxanthin in *C. reinhardtii*. Blue light exposure causes 1.62 times enhanced accumulation when compared to red light. We reasoned that astaxanthin is produced at the diploid zygospore stage in *C. reinhardtii* [34] and blue light regulates the life cycle of *C. reinhardtii* [35]. It was shown that reduced phototropin in *C. reinhardtii* leads to impairment of life cycle mainly at gametogenesis stage, mating ability and zygotes germination. Thus, it suggested that modulation of light conditions resulted in astaxanthin accumulation in *C. reinhardtii* without genetic modification. In *Haematococcus pluvialis*, commercial strain for astaxanthin production, astaxanthin accumulation is linked to phototropin [36]. This also indicates that the astaxanthin accumulation is phototropin networking modulated. To confirm this, we investigated whether carotenoid metabolite accumulation particularly, astaxanthin gets affected by the disrupted phototropin. For this, we used CRISPR-Cas knock out mutant of phototropin. We determined the expression of important rate limiting genes (phytoene synthase and β-carotene hydroxylase) of astaxanthin metabolism [37]. Our data suggested that these genes were significantly downregulated in phot mutants (Figure 3B, S2F). Notably, Ct value of β-carotene hydroxylase in phot mutant was around 36-37. While Ct value for β-carotene ketolase was undetermined. Further, we observed that the carotenoid content and chlorophyll content were reduced under the influence of blue and red light in the phot mutant CC-125 Δ phot-B5 (Figure 3C), Additional data 3). A substantial 2.5-fold and 3.8-fold decrease in carotenoid content was observed in the phot mutants, respectively, under blue and red light illumination, as compared to the wild type. A significant 64 % reduction in astaxanthin content was noted in phot mutant when compared with wild type (Figure 3D, S2E). This suggests that phototropin is the key mediator in carotenoid metabolism [6]. It would be interesting to see whether other photoreceptor also regulates astaxanthin accumulation.

Further, we performed qRT-PCR to detect the potency of anti-inflammatory effects due to astaxanthin in cell lines. Concentrations showing minimal cytotoxicity and ≥80% cell viability were selected for subsequent qRT-PCR and anti-inflammatory studies (Figure S5). We used actin as reference gene to normalize the expression. The result demonstrated that the *C. reinhardtii*, when exposed to blue light treatment under specific conditions (photoperiod of 12 hours light and 12 hours dark with a photon flux density (PFD) of 120 µmol m⁻² s⁻¹ continuous 3-4 days), resulted in downregulation of inflammatory genes (Figure S6). This suggested that the astaxanthin obtained has anti-inflammatory activity.

### 3.5 Quantitative proteomics revealed the phototropin drives upregulation of metabolite content in *C. reinhardtii*

Further, the comparative whole cell proteomics was performed to unravel the key enzymes and molecules playing a role in the upregulation of specific metabolite production under different light regimes. The data revealed a significant impact of blue and red light on the whole cell proteome of *C. reinhardtii* quantitatively. A total of 4621 proteins were identified, and their expression profiles in blue vs. red light conditions are represented by a heat map (Figure 4A). A list of differentially expressed proteins (DEPs) in *C. reinhardtii* strain CC-125 in response to blue vs. red light is given in the supporting information (Table S1). The upregulated proteins, both in blue and red-light conditions, are shown in the volcano plot (Figure 4B). Further, when both the light conditions were compared, out of the total proteins identified by the label-free proteomics, 265 proteins showed significant changes in their expression (Figure 4C). Among them, the key enzymes of the carotenoid metabolism such as phytoene synthase, phytoene desaturase, 15-cis-phytoene synthase, carotenoid 9,10-dioxygenase, zeaxanthin epoxidase, zeta-carotene desaturase, lycopene beta-cyclase etc., was found to be significantly upregulated under blue light. The carotenoid qRT-PCR also suggested that phytoene synthase, phytoene desaturase and zeaxanthin epoxidase revealed a higher expression only in response to blue light, whereas reduced expression was observed in red light (Figure 4D). DEPs mainly belonging to photosynthesis, photoprotection, and carotenoid metabolism were upregulated in blue light vs. red light. Among photoreceptors, phototropin (log 2-fold-0.5, p value 0.2) remains significantly unaffected when compared between red and blue light. Cryptochrome and Cryptochrome-DASH was upregulated while archaeal-type opsin was downregulated. Further, the principal component analysis and correlation coefficient show the difference and similarity between the proteome at different light conditions (Figure S7). Thus, this indicates imperative effect of light on algal physiology. Apart from these, the prolonged diurnal blue light exposure leads to abundance of proteomes for protein kinases / phosphatases (Serine-threonine kinases, MAPK-like kinases, Casein kinase / CDPK-type proteins), calcium signalling components (Ca²⁺ channels, Calmodulin-like proteins,) Chloroplast redox sensors, ROS-responsive regulators, Tetrapyrrole / plastid stress signalling proteins, transcriptional regulators like bZIP / MYB-like TFs, light-responsive TFs (ELONGATED HYPOCOTYL-like, CONSTANS-like analogs in algae), stress-integrating regulators (NAC-like, AP2-likeetc. These suggest that Phototropin-modulates functionally clustered carotenoid biosynthetic gene modules signalling. Interestingly, our data also showed zygote-specific Zys3-like protein high abundance under the influence of blue light, which is responsible for Further, the GO enrichment analysis was performed to see the over-representation of different pathways. In comparison to red light, the blue light has higher impact on the DEPs, of secondary metabolic processes (Figure S8). The list GO categories enriched among significantly differentially expressed proteins is shown in the supplementary file (Table S2, Additional data 5). Figure S8 is a representation of the top 20 GO terms of DEP in BL vs those in RL. The KEGG pathway analysis was done for the significantly upregulated DEP. Proteins belonging to photosynthesis, carbon metabolism, secondary metabolite production were among top 20 GO terms.

**Figure 4:**
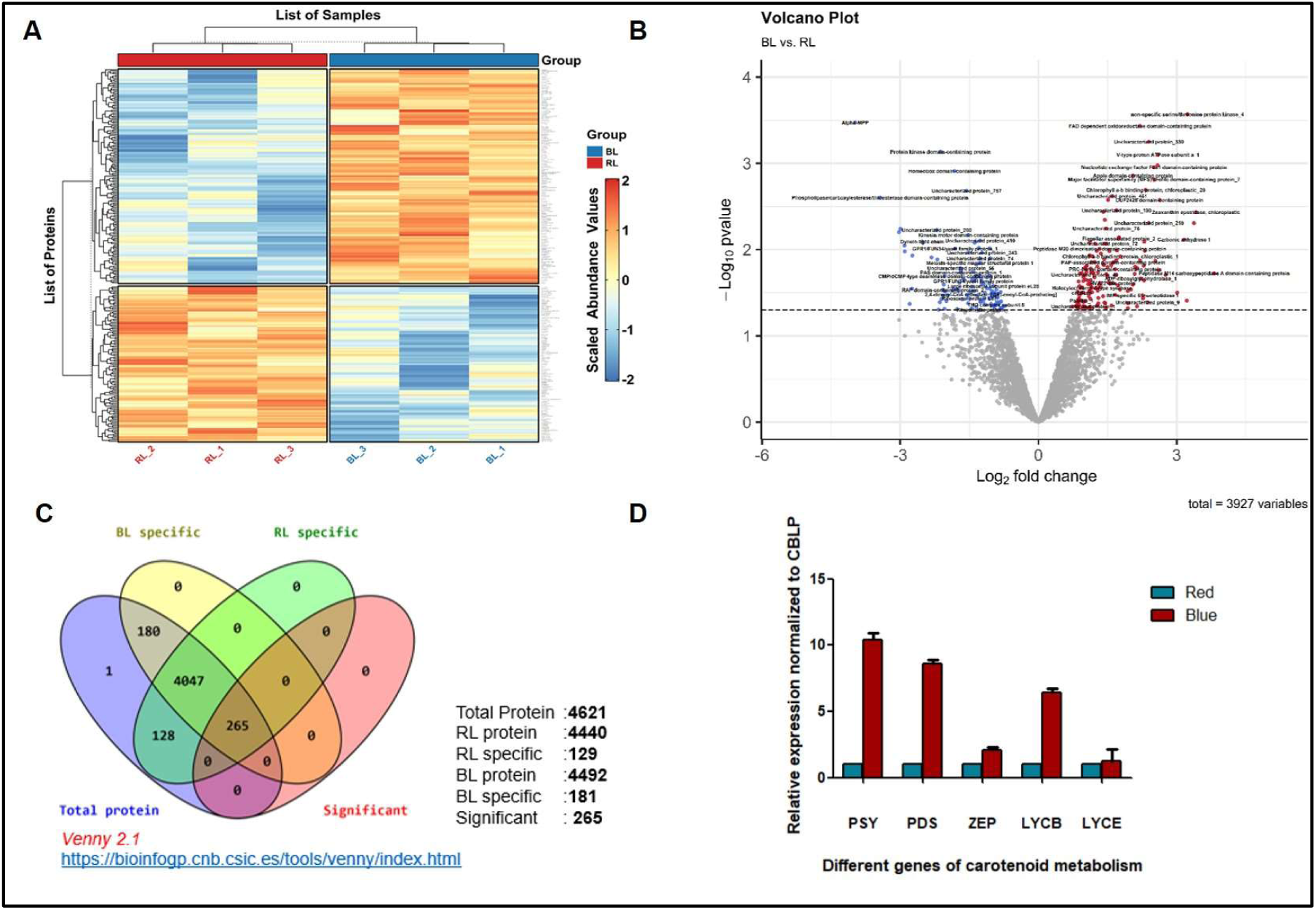
Comparative proteomics of *C. reinhardtii* illuminated with blue (BL) and red light (RL). (A) The heat map for the DEPs for BL vs. RL. (B)Volcano plot showing proteins significantly downregulated (blue colour) and upregulated (red colour) for DEPs obtained for BL vs. RL. (C) Represents a Venn diagram for the distribution of protein observed under BL and RL. (D) Represents the expression profile of different genes like *ZEP, PSY, PDS, LYCB and LYCE* in *C. reinhardtii*, after blue light and red-light treatment (Control samples-sample exposed to blue light). Values are mean ± SE (n = 2, technical replicates and 2 biological replicates). PSY; Phytoene synthase, LYCB; lycopene β-cyclase, ZEP; zeaxanthin epoxidase, PDS; phytoene desaturase, and LYCE; lycopene ε-cyclase.

### 3.6 Light driven biosynthetic gene clusters (BGC)-mediated metabolite production in *C. reinhardtii*

Additionally, we also examined the role of light in modulating biosynthetic gene clusters in opto-biomanufacturing. The enzymes and molecules pertaining to these pathways are encoded in biosynthetic gene clusters [13]. BGCs in bacterial and fungal genomes are frequently investigated for a detailed understanding of metabolite accumulation. Numerous BGCs have been identified in plants [38,39,40]. Limited information is available for algal BGCs. The research for algal BGCs is mainly centralised towards the identification of BGCs [41,42]. In our study, for the first time, we established light-regulated BGCs-mediated biomolecule production in green algae. Based on our experimental and systems biology-based data, it is postulated that the molecular components of biosynthetic gene clusters accountable for specific metabolite production would also be influenced by different illumination conditions perceived by the native sensory photoreceptor(s). If that’s the case, it would enhance the accumulation of a specific metabolite and regulation of relevant signalling pathways under carefully controlled light conditions (quality, quantity, duration, pattern and their synchronization). In light of this discussion, we have therefore identified the BGCs and performed in-depth analysis by integrating our proteomics data and systems biology in specific light conditions. Interestingly, BGC core component, fatty acid desaturase and 3-oxoacyl-[acyl-carrier-protein] synthase (shows interaction with phytoene synthase which is directly regulated by phototropin), were upregulated. Remarkably, the molecular components of cluster 3, starch synthase and beta-carotene hydroxylase, were among the DEPs obtained from our proteomics data. We also noticed negligible expression of *beta-carotene hydroxylase* in phot mutant, suggesting role of photoreceptor controlled BGC-mediated in channelising the metabolic flux towards β-carotene and ketocarotenoid production. It is to note that, the important enzyme for lipid metabolism, fatty acid desaturase, was found to be upregulated under blue light illumination in DEP. Similarly, identified BGC clusters comprise genes for important steps in glycosylation (glycosyl-transferase 1). Our proteomics data also reported upregulation of glycosyl transferase family protein (see supplementay file table S1 and S2). Likewise, the gene cluster also showed interaction with the enzymes mediating lipids, mainly triacylglycerol metabolism, an important component of biofuel production. This strengthens our concept of light-driven BGCs mediated metabolite production in microalgae. Moreover, the molecular component of *K. nitens* (terrestrial green alga) BGCs cluster showed direct interaction with different photoreceptor(s) (rhodopsin, phototropin and cryptochrome). This provides a basis for photoreceptor-guided BGC-mediated opto-biomanufacturing of valuable metabolites in the green lineage (Figure 5).

**Figure 5:**
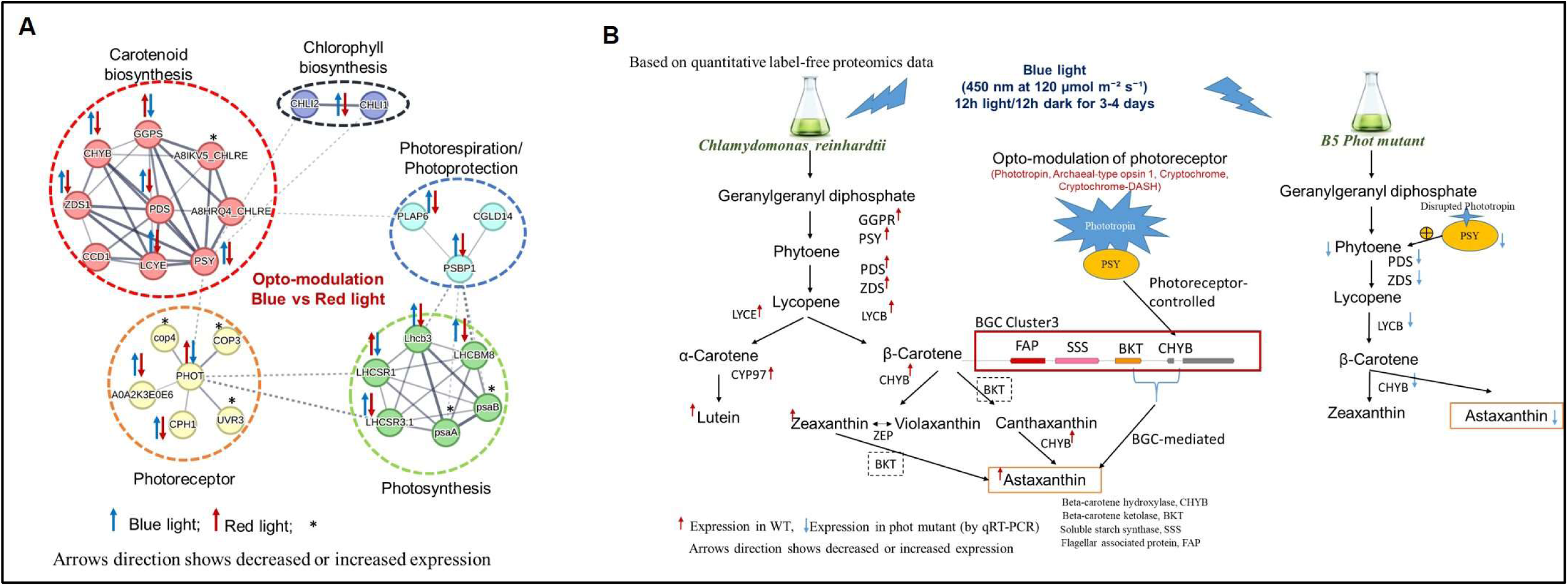
**Proposed mechanistic basis for phototropin modulated BGC-mediated metabolite production in *C. reinhardtii.*** (A): Represents the major pathways affected. (B): Represents the schematic representation of phototropin-modulated BGC-mediated metabolite production in *C. reinhardtii.* PSY; Phytoene synthase, CHYB; beta-carotene hydroxylase, LYCB; lycopene β-cyclase, ZEP; zeaxanthin epoxidase, PDS; phytoene desaturase, and Bp450; P450 cytochrome–dependent β-ring carotene hydroxylase genes.

Additionally, we also export the BGC concept in higher plants. BGCs related to terpenoid metabolism are well identified and characterized in rice and *Arabidopsis*. Most of the BGCs in rice (*O. sativa* Indica) and *A. thaliana* belongs to terpene and saccharides. Similarly, in *K. nitens* (closely related to higher plants) among 7 identified BGCs, 5 are for saccharide, one for polyketide and one for lignin. The core domain of these clusters was similar to that of rice and *Arabidopsis* (the data for BGCs for rice and *Arabidopsis* is available under pre-calculated results in plantiSMASH database). The bio-curated PPI of photoreceptors and selected BGCs of *A. thaliana* and rice showed crosstalk with each other (Figure S9 and S10). Several studies established illumination impact on flavonoid and terpenoid biosynthetic pathway in rice [43,44,45,46]. Hence, photoreceptor modulation can be employed for development of stress resilient smart crop [47]. Numerous published literatures suggest light-regulated metabolite production and signalling in higher plants. Thus, it is suggested that opto-modulation of BGCs holds potential for manufacturing of important commercial bioproducts and open up a new biotechnological avenue for biomanufacturing of valuable products from green lineage.

## 4 Discussion

Light, being an important environmental cue, not only drives photosynthesis, but it also spatiotemporally regulates photosynthesis machinery [3,4,5,6,7]. Earlier results demonstrated that low fluence blue light in coordination with red light affects the carotenoid and chlorophyll biosynthesis [6]. Our result, for the first time, showed insight into the comparative molecular components when algal cells were grown under specific light quality (blue and red light). The comparative quantitative label-free proteomics data showed that enzymes and proteins involved in photosynthesis and chlorophyll metabolism were upregulated in the presence of blue light as compared to red light (Figure 4). On the contrary, ribulose bisphosphate carboxylase/oxygenase and starch synthase were upregulated by red light. Similarly, Li et al. [9] also showed 38% increased biomass under red light illumination as compared to blue light. Thus, it suggested that although red light illumination positively affects the algal biomass, blue light illumination can significantly accumulate more metabolites at low biomass. Short blue light pulses rapidly upregulate transcripts for phytoene desaturase (PDS) and phytoene synthase, key rate-limiting enzymes in carotenoid biosynthesis, via PHOT signaling. This leads to transient increases in carotenoids like β-carotene and zeaxanthin for photoprotection, but levels normalize post-exposure without entrainment. Extended 12 h blue/12 h dark cycles synchronize carotenoid gene expression diurnally, sustaining higher PDS and related transcript levels for ongoing biosynthesis synchronized with photosynthesis. Additionally, the downstream enzymes and molecules of carotenoid metabolism (phytoene synthase, phytoene desaturase, phytoene dehydrogenase, carotenoid 9,10-dioxygenase, lycopene beta-cyclase, zeta-carotene desaturase and zeaxanthin epoxidase) were upregulated under blue light illumination with respect to red light (as shown in the supplementary table S1 and S2). However, most of the enzymes associated with primary metabolism remain unaffected in response to blue and red-light illumination. Interestingly, we also found light-driven accumulation of astaxanthin without genetic modification. Our data suggests that photoreceptor networking particularly, phototropin coordinates astaxanthin accumulation. Although the differential expression of enzymes β-carotene ketolase specific for conversion of β-carotene to canthaxanthin (an intermediate) were not found through proteomics data. While β-carotene hydroxylase converting canthaxanthin to astaxanthin were upregulated (Figure 3). This is because we have done comparative quantitative label-free proteomics between red and blue light. It would be interesting if such studies will be done comparing white light with other light conditions, at different developmental stages [32,33]. Notably, both these enzymes were obtained as molecular components of one of the identified BGCs in *C. reinhardtii.* Additionally, our proteomics data also showed zygote-specific Zys3-like protein high abundance under the influence of blue light. The systems biology-based computational analysis and experimental data in our study elicit that the biosynthetic pathway is regulated by the coordinated interactions of photoreceptors. Our study revealed interaction of photoreceptor(s) with the protein family of carotenoid and chlorophyll pathway, cell signalling and important metabolic pathways (Figure 1). The biochemical assay and qRT-PCR analysis of *phot* mutant showed that disruption of *photo* gene resulted in insignificant expression of *BKT* and *CHYB* gene and astaxanthin accumulation. Recent study also established the role of UVR8 photoreceptor in flavonoid metabolism. First-ever, we are providing biochemical, genetic, transcriptomic, quantitative proteomics and systems biology evidences for opto-biomanufacturing of bioactive from green lineage via modulation of phototropin network with artificial illumination (without any genetic modification). This suggested that artificial modulation of algal photoreceptor(s) can regulate biomass and improve their biochemical makeup for biorefinery of high-value bioproducts. High abundance of BGC core components under blue light exposure pave way for establishment of opto-biomanufacturing strategies to generate bioactive without genetic modification through cascade biorefinery.

## 5 Conclusions

In recent years, genetic engineering is mainly driving the research concerning carotenoid metabolism and enhanced metabolites accumulation in *C. reinhardtii*. The present study showed the imperative effect of light-regulated phototropin networking with BGCs on the biomanufacturing of valuable metabolites in green algae *C. reinhardtii*. By fine tuning the light conditions astaxanthin biomanufacturing was observed by simply modulating light conditions in terms of quality (red and blue), intensity (120 µmol m⁻² s⁻¹), and duration (continuous exposure with 12-hour light/12-hour dark phototropism for 3-4 days). Moreover, we have offered a new aspect of opto-modulated biosynthetic gene cluster-mediated metabolite production in green lineage specifically in *C. reinhardtii* (aquatic alga), *K. nitens*, a green terrestrial alga having an evolutionary link between land and aquatic plants, *A. thaliana* and *O. sativa* (higher plants). This approach could also be employed for the sustainable biomanufacturing of specific metabolites simply by modulating illumination conditions not only in microalgae but also in the green lineage system. However, in-depth research is required in this area. Overall, our result suggests a crucial role of illumination conditions in the biomanufacturing of sustainable and safe metabolite production from microalgae.

## Author Contributions

A.S. and S.K. conceived the project. R.S. designed the experiments in the guidance of S.K. and A.S. and drafted the manuscript with the help of A.S., H.A. and S.K. H.A. performed growth and maintenance of algal culture under different conditions, growth curve analysis and cell morphology visualization experiments for the publication. R.S. conducted immunoblotting, qRT-PCR, and HPLC, bioinformatics, data curation and analysis. All authors reviewed and approved the final manuscript. N. performed the bioassay to test the potency of anti-inflammatory effect of algal extract in cell line.

## Author’s statement

All authors have seen and approved the manuscript. This work is original and not under consideration or published anywhere.

## Acknowledgements

S.K and A.S. are highly thankful to the DBT project grants BT/PR53866/BSA/33/107/2024 and BT/PR34567/BRB/10/1822/2019 for providing financial assistance. RS acknowledge DBT-RA program (DBT-RA/2022/July/N/2560). H.A. acknowledge DBT (BT/PR34567/BRB/10/1822/2019) for providing Manpower.

**Appendix:** Supplementary information and Additional data 1-6

## Abbreviations

Chl a: Chlorophyll a
Chl b: Chlorophyll b
TAP: Tris-Acetate-Phosphate
PFD: Photon Flux Density
PBS: Phosphate-Buffered Saline
PFA: Paraformaldehyde
HPLC-MS/MS: High-Performance Liquid Chromatography coupled with Tandem Mass Spectrometry
MS: Mass Spectrometry
plantiSMASH: Plant Specialized Metabolite Analysis

## Glossary

Photosynthesis: The physio-chemical process converting light energy to chemical energy utilising carbon dioxide and water and releasing molecular oxygen as a byproduct.
Photoreceptors: Specialized cells or protein molecules that detect light and convert it into a downstream signal allowing organisms to perceive and respond to their light environment.
Label-Free Comparative Quantitative Proteomics: A technique used to determine the relative or absolute amounts of proteins in a biological sample, by comparing protein abundance between different conditions.
Biosynthetic Gene Cluster: The biosynthetic gene clusters (BGCs) are collections of interconnected genes, including operons and independently transcribed genes that collectively carry out the production of certain natural products.
Opto-modulation: The application of light to regulate specific physiological processes in an organism by activating or deactivating relevant cellular processes.
Opto-Biomanufacturing: The use of light or light based tools to control biological systems for producing the desired product.
Protein-Protein Interactions Network: The computational prediction and visualization of protein interactions, to infer functional associations.
Gene Ontology: A comprehensive, hierarchically structured classification system that describes the functions of genes and proteins across all organisms, categorizing them into three main aspects: molecular function, cellular component, and biological process.
Photon Flux Density: A measure of the number of photons incident on a given surface area per unit of time to quantify the amount of light available for photosynthesis.

