## Supplementary materials for "Phototropin modulates biosynthetic gene clusters network mediated biomanufacturing of anti-inflammatory metabolites in green lineage simply by illumination"

*To whom correspondence should be addressed.

†ORCID identifiers: R.S. 0000-0002-1520-2794; H.A. 0009-0005-8663-5502; A.S. 0000-0002-3568-1569; P.H. [0000-0003-3589-6452](https://orcid.org/0000-0003-3589-6452); S.K. 0000-0001-5428-4297

**Supplementary Tables**

**Table S1:** **List of differentially expressed proteins (DEPs) in *C. reinhardtii* strain CC-125 in response to Blue vs. Red light obtain from quantitative label-free proteomics.**

| ***Accession*** | ***Description*** |
| --- | --- |
| A0A250WSQ8 | *Uncharacterized protein* |
| A0A250X9Z6 | *Protein kinase domain-containing protein* |
| A0A250XIC7 | *Chlorophyll a-b binding protein, chloroplastic* |
| A0A250XIW1 | *Dynein light chain* |
| A0A250XNL3 | *Ribosomal protein L37* |
| A0A2K3CNG5 | *Uncharacterized protein_1* |
| A0A2K3CNN3 | *HVA22-like protein* |
| A0A2K3CP17 | *GPR1/FUN34/yaaH family protein* |
| A0A2K3CP19 | *GPR1/FUN34/yaaH family protein_1* |
| A0A2K3CPA3 | *THO complex subunit 5* |
| A0A2K3CQA0 | *PAP-associated domain-containing protein* |
| A0A2K3CQB5 | *Alpha-type protein kinase domain-containing protein* |
| A0A2K3CQS1 | *Peptidase M20 dimerisation domain-containing protein* |
| A0A2K3CR15 | *2,4-dienoyl-CoA reductase [(3E)-enoyl-CoA-producing]* |
| A0A2K3CR95 | *RAP domain-containing protein* |
| A0A2K3CRG7 | *Meiosis-specific nuclear structural protein 1* |
| A0A2K3CRJ5 | *PRC-barrel domain-containing protein* |
| A0A2K3CRT1 | *Kinesin motor domain-containing protein* |
| A0A2K3CSH8 | *Uncharacterized protein_2* |
| A0A2K3CTA4 | *Vacuolar import/degradation Vid27 C-terminal domain-containing protein* |
| A0A2K3CTF1 | *RRM domain-containing protein* |
| A0A2K3CTG0 | *Uncharacterized protein_3* |
| A0A2K3CU79 | *Holocytochrome c-type synthase* |
| A0A2K3CUF3 | *Flagellar associated protein* |
| A0A2K3CUS3 | *Peptidase S8/S53 domain-containing protein* |
| A0A2K3CUT5 | *Uncharacterized protein_4* |
| A0A2K3CV53 | *Uncharacterized protein_5* |
| A0A2K3CVC3 | *Methylenetetrahydrofolate reductase (NAD(P)H)* |
| A0A2K3CVD3 | *AAA+ ATPase domain-containing protein* |
| A0A2K3CVF0 | *peptidylprolyl isomerase* |
| A0A2K3CVV7 | *Palmitoyl-protein thioesterase 1* |
| A0A2K3CW67 | *Translation initiation factor eIF2B subunit alpha* |
| A0A2K3CWJ2 | *Uncharacterized protein_6* |
| A0A2K3CWP8 | *RAP domain-containing protein_1* |
| A0A2K3CWX3 | *DUF2428 domain-containing protein* |
| A0A2K3CX72 | *Uncharacterized protein_7* |
| A0A2K3CX83 | *Uncharacterized protein_8* |
| A0A2K3CXH6 | *Flagellar associated protein_1* |
| A0A2K3CXR7 | *ABC transporter domain-containing protein* |
| A0A2K3CYN7 | *RING-type domain-containing protein* |
| A0A2K3CYQ5 | *3-hydroxyisobutyrate dehydrogenase* |
| A0A2K3CZM2 | *CMP/dCMP-type deaminase domain-containing protein* |
| A0A2K3D0C5 | *non-specific serine/threonine protein kinase* |
| A0A2K3D1V1 | *Uncharacterized protein_9* |
| A0A2K3D1W6 | *FAS1 domain-containing protein* |
| A0A2K3D2W6 | *Uncharacterized protein_10* |
| A0A2K3D2Z9 | *1-alkyl-2-acetylglycerophosphocholine esterase* |
| A0A2K3D379 | *TRAM domain-containing protein* |
| A0A2K3D385 | *Aldehyde dehydrogenase domain-containing protein* |
| A0A2K3D3N5 | *SF4 helicase domain-containing protein* |
| A0A2K3D3S9 | *Uncharacterized protein_11* |
| A0A2K3D468 | *Kinesin-like protein* |
| A0A2K3D4K9 | *Transcription initiation factor TFIID subunit 5* |
| A0A2K3D581 | *2-oxoglutarate dehydrogenase, mitochondrial* |
| A0A2K3D7C4 | *Large ribosomal subunit protein eL28* |
| A0A2K3D7S2 | *V-type proton ATPase subunit a* |
| A0A2K3D836 | *Uncharacterized protein_12* |
| A0A2K3D937 | *SAM domain-containing protein* |
| A0A2K3D996 | *Hexosyltransferase* |
| A0A2K3D9C0 | *Peptidase S9 prolyl oligopeptidase catalytic domain-containing protein* |
| A0A2K3D9G6 | *Nuclear pore complex protein Nup85* |
| A0A2K3D9I3 | *Uncharacterized protein_13* |
| A0A2K3D9M9 | *Uncharacterized protein_14* |
| A0A2K3DA34 | *Ion transport domain-containing protein* |
| A0A2K3DAI3 | *HTH La-type RNA-binding domain-containing protein* |
| A0A2K3DAK6 | *Pre-rRNA-processing protein Ipi1 N-terminal domain-containing protein* |
| A0A2K3DAQ7 | *Peptidase S49 domain-containing protein* |
| A0A2K3DAY3 | *Mitochondrial glycoprotein* |
| A0A2K3DB02 | *DAGKc domain-containing protein* |
| A0A2K3DB58 | *C2 domain-containing protein* |
| A0A2K3DCA6 | *peptidylprolyl isomerase_1* |
| A0A2K3DCG6 | *Uncharacterized protein_15* |
| A0A2K3DD56 | *Uncharacterized protein_16* |
| A0A2K3DD93 | *ADP/ATP translocase* |
| A0A2K3DDB9 | *FAS1 domain-containing protein_1* |
| A0A2K3DDD4 | *Uncharacterized protein_17* |
| A0A2K3DDQ3 | *Tbc2 translation factor, chloroplastic* |
| A0A2K3DEI2 | *VTT domain-containing protein* |
| A0A2K3DEQ0 | *Nucleotide exchange factor Fes1 domain-containing protein* |
| A0A2K3DER4 | *Apple domain-containing protein* |
| A0A2K3DF23 | *STAS domain-containing protein* |
| A0A2K3DF70 | *Uncharacterized protein_18* |
| A0A2K3DFE4 | *Uncharacterized protein_19* |
| A0A2K3DGK9 | *PAS domain-containing protein* |
| A0A2K3DGS4 | *VWFA domain-containing protein* |
| A0A2K3DGU2 | *Assimilatory sulfite reductase (ferredoxin)* |
| A0A2K3DGY3 | *Uncharacterized protein_20* |
| A0A2K3DHY3 | *Major facilitator superfamily (MFS) profile domain-containing protein* |
| A0A2K3DI85 | *U-box domain-containing protein* |
| A0A2K3DIA5 | *GB1/RHD3-type G domain-containing protein* |
| A0A2K3DIV5 | *Protein transport protein Sec61 subunit beta* |
| A0A2K3DJ31 | *Activator of Hsp90 ATPase AHSA1-like N-terminal domain-containing protein* |
| A0A2K3DL29 | *EF-hand domain-containing protein* |
| A0A2K3DLQ3 | *Cytochrome P450* |
| A0A2K3DMY1 | *FAD dependent oxidoreductase domain-containing protein* |
| A0A2K3DN52 | *Uncharacterized protein_21* |
| A0A2K3DNQ7 | *Dethiobiotin synthase* |
| A0A2K3DPT9 | *EF-hand domain-containing protein_1* |
| A0A2K3DPU1 | *Flagellar associated protein_2* |
| A0A2K3DQ06 | *Uncharacterized protein_22* |
| A0A2K3DQ47 | *Methyltransferase type 11 domain-containing protein* |
| A0A2K3DQS7 | *Aminotransferase-like plant mobile domain-containing protein* |
| A0A2K3DQU3 | *Pherophorin domain-containing protein* |
| A0A2K3DQY4 | *Uncharacterized protein_23* |
| A0A2K3DRH6 | *Uncharacterized protein_24* |
| A0A2K3DRW2 | *Uncharacterized protein_25* |
| A0A2K3DSN9 | *AAA+ ATPase domain-containing protein_1* |
| A0A2K3DTU2 | *polyribonucleotide nucleotidyltransferase* |
| A0A2K3DTY7 | *Uncharacterized protein_26* |
| A0A2K3DUA0 | *OCRE domain-containing protein* |
| A0A2K3DUD4 | *Surfeit locus 2* |
| A0A2K3DUK8 | *IMP-specific 5'-nucleotidase 1* |
| A0A2K3DV09 | *Sulfhydryl oxidase* |
| A0A2K3DVA7 | *tyrosine--tRNA ligase* |
| A0A2K3DVJ8 | *Uncharacterized protein_27* |
| A0A2K3DVQ5 | *Uncharacterized protein_28* |
| A0A2K3DWB3 | *Elongation factor Tu, chloroplastic* |
| A0A2K3DWL6 | *Phospholipase/carboxylesterase/thioesterase domain-containing protein* |
| A0A2K3DWN6 | *Ubiquitin-like modifier-activating enzyme ATG7* |
| A0A2K3DWV7 | *Bromo domain-containing protein* |
| A0A2K3DX15 | *Uncharacterized protein_29* |
| A0A2K3DXB3 | *Homeobox domain-containing protein* |
| A0A2K3DXH1 | *Uncharacterized protein_30* |
| A0A2K3DXT5 | *methylcrotonoyl-CoA carboxylase* |
| A0A2K3DYJ1 | *Protein NEOXANTHIN-DEFICIENT 1* |
| A0A2K3DYL4 | *Patatin* |
| A0A2K3DYS1 | *Uncharacterized protein_31* |
| A0A2K3DZJ4 | *Mitochondrial carrier protein* |
| A0A2K3E146 | *SnoaL-like domain-containing protein* |
| A0A2K3E1M2 | *Uncharacterized protein_32* |
| A0A2K3E2K9 | *Histone H2A/H2B/H3 domain-containing protein* |
| A0A2K3E2S5 | *RAP domain-containing protein_2* |
| A0A2K3E2Y6 | *C2 domain-containing protein_1* |
| A0A2K3E2Z8 | *K Homology domain-containing protein* |
| A0A2K3E338 | *SBP-type domain-containing protein* |
| A0A2K3E3E1 | *Glycoside hydrolase family 42 N-terminal domain-containing protein* |
| A0A2K3E4Q3 | *Uncharacterized protein_33* |
| A0A2K3E4V9 | *Acid phosphatase* |
| A0A2K3E4W2 | *Rhodanese domain-containing protein* |
| A0A2K3E592 | *Cytosol aminopeptidase domain-containing protein* |
| A0A2K3E5B0 | *N-acetyltransferase domain-containing protein* |
| A0A2K3E5E3 | *t-SNARE coiled-coil homology domain-containing protein* |
| A0A2K3E5Q7 | *Tocopherol cyclase* |
| A0A2K3E5T3 | *ADP-ribosylglycohydrolase* |
| A0A2K3E6L6 | *Rhodanese domain-containing protein_1* |
| A0A2K3E6L7 | *adenylate kinase* |
| A0A2K3E6S5 | *PDZ domain-containing protein* |
| A0A2K3E7K0 | *Uncharacterized protein_34* |
| A0A2K3E861 | *CobW C-terminal domain-containing protein* |
| A0A2K3E8F4 | *DJ-1/PfpI domain-containing protein* |
| A0A2K3E8I4 | *CCAAT-binding factor domain-containing protein* |
| A0A2K3E8L0 | *DNA-directed RNA polymerase I subunit rpa49* |
| A0A387J5P9 | *Photosystem I P700 chlorophyll a apoprotein A1 (Fragment)* |
| A0A6T8Q0U8 | *Uncharacterized protein_35* |
| A0A7R9V082 | *ADP/ATP translocase_1* |
| A0A7R9VQU9 | *catalase* |
| A0A7S0R5D6 | *Aspartokinase* |
| A0A7S0RS96 | *non-specific serine/threonine protein kinase_1* |
| A0A7S0RY46 | *PAS domain-containing protein_1* |
| A0A7S0X021 | *CCR4-NOT transcription complex subunit 1* |
| A0A835S917 | *Protein kinase domain-containing protein_1* |
| A0A835SC30 | *PHD-type domain-containing protein* |
| A0A835SCK1 | *Nitric oxide synthase-interacting protein zinc-finger domain-containing protein* |
| A0A835SHM2 | *Uncharacterized protein_36* |
| A0A835SI34 | *Uncharacterized protein_37* |
| A0A835SJE1 | *formate C-acetyltransferase* |
| A0A835SKV8 | *Uncharacterized protein_38* |
| A0A835SLH6 | *C2H2-type domain-containing protein* |
| A0A835SS52 | *Protein kinase domain-containing protein_2* |
| A0A835STK4 | *Uncharacterized protein_39* |
| A0A835SUV8 | *3-methyl-2-oxobutanoate dehydrogenase (2-methylpropanoyl-transferring)* |
| A0A835SVJ6 | *Methyltransferase type 11 domain-containing protein_1* |
| A0A835T1N5 | *Paladin* |
| A0A835T2M4 | *Alpha-MPP* |
| A0A835T5B7 | *ShKT domain-containing protein* |
| A0A835T6E5 | *C-CAP/cofactor C-like domain-containing protein* |
| A0A835TAF7 | *Zeaxanthin epoxidase, chloroplastic* |
| A0A835TAJ4 | *Protein kinase domain-containing protein_3* |
| A0A835TEY3 | *COP9 signalosome complex subunit 6* |
| A0A835TF64 | *ABC transmembrane type-1 domain-containing protein* |
| A0A835TM17 | *WPP domain-containing protein* |
| A0A835TRP3 | *Protein kinase domain-containing protein_4* |
| A0A835VS90 | *Uncharacterized protein_40* |
| A0A835W2W8 | *Serine/threonine-protein phosphatase* |
| A0A835W3C9 | *Signal recognition particle subunit SRP68* |
| A0A835WC61 | *GPI inositol-deacylase* |
| A0A835WDQ4 | *ABC transporter domain-containing protein_1* |
| A0A835WFF9 | *SRP9 domain-containing protein* |
| A0A835WH08 | *Band 7 domain-containing protein* |
| A0A835WIM5 | *U-box domain-containing protein_1* |
| A0A835WKJ6 | *Apple domain-containing protein_1* |
| A0A835WM92 | *Clusterin-associated protein 1* |
| A0A835WPC0 | *Uncharacterized protein_41* |
| A0A835WQ96 | *Uncharacterized protein_42* |
| A0A835WQH2 | *SOUL heme-binding protein* |
| A0A835WRP4 | *Altered inheritance of mitochondria protein 24, mitochondrial* |
| A0A835WTA8 | *PH domain-containing protein* |
| A0A835WTP2 | *L-aspartate oxidase* |
| A0A835WTQ3 | *Uncharacterized protein_43* |
| A0A835WTX7 | *Glutathione S-transferase* |
| A0A835WUY6 | *Fungal lipase-like domain-containing protein* |
| A0A836B1J3 | *Activating signal cointegrator 1 complex subunit 3* |
| A0A836B653 | *5-methyltetrahydropteroyltriglutamate--homocysteine S-methyltransferase* |
| A0A836B8M7 | *Uncharacterized protein_44* |
| A0A836B9C3 | *Fatty acid hydroxylase domain-containing protein* |
| A0A836B9K3 | *Uncharacterized protein_45* |
| A0A836BCY4 | *Uncharacterized protein_46* |
| A0FK50 | *Chloroplast Rat1* |
| A8HNJ6 | *Ubiquinol-cytochrome c chaperone domain-containing protein* |
| A8HNU9 | *Uncharacterized protein_47* |
| A8HP55 | *Large ribosomal subunit protein uL18 C-terminal eukaryotes domain-containing protein* |
| A8HPY4 | *Uncharacterized protein_48* |
| A8HW77 | *CBM20 domain-containing protein* |
| A8HXE1 | *Uncharacterized protein_49* |
| A8HZ72 | *PsbP C-terminal domain-containing protein* |
| A8HZF9 | *Flagellar associated protein_3* |
| A8I594 | *Uncharacterized protein_50* |
| A8I7P5 | *magnesium chelatase* |
| A8IAD4 | *SAM domain-containing protein_1* |
| A8IC56 | *Uncharacterized protein_51* |
| A8IE17 | *Uncharacterized protein_52* |
| A8ILA3 | *Arf* |
| A8INE5 | *Peptidyl-prolyl cis-trans isomerase CYP38-like PsbQ-like domain-containing protein* |
| A8IQ05 | *Succinate--CoA ligase [ADP-forming] subunit beta, mitochondrial* |
| A8IQG4 | *GPR1/FUN34/yaaH family protein_2* |
| A8ISV8 | *Uncharacterized protein_53* |
| A8IUB5 | *Arsenate reductase* |
| A8IUR4 | *EndoU domain-containing protein* |
| A8IVP1 | *Uncharacterized protein_54* |
| A8IX19 | *Single-stranded DNA binding protein Ssb-like OB fold domain-containing protein* |
| A8IXZ2 | *UspA domain-containing protein* |
| A8J071 | *Protein BIG1* |
| A8J0B1 | *CBM20 domain-containing protein_1* |
| A8J0U5 | *amino-acid N-acetyltransferase* |
| A8J1M7 | *HNH nuclease domain-containing protein* |
| A8J1Y2 | *Carboxypeptidase* |
| A8J940 | *Uncharacterized protein_55* |
| A8J9T5 | *Thiamine thiazole synthase, chloroplastic* |
| A8JA42 | *Intraflagellar transport protein 56* |
| A8JB30 | *Peptidase M14 carboxypeptidase A domain-containing protein* |
| A8JDA2 | *Thioredoxin domain-containing protein* |
| A8JDW2 | *TPM domain-containing protein* |
| A8JEP4 | *Uncharacterized protein_56* |
| A8JFF7 | *GDT1 family protein* |
| A8JHG4 | *Cyanobacterial aminoacyl-tRNA synthetase CAAD domain-containing protein* |
| A8JIB8 | *Uncharacterized protein_57* |
| D2K6F1 | *Sodium/sulfate cotransporter 2* |
| P05725 | *DNA endonuclease I-CreI* |
| P14273 | *Chlorophyll a-b binding protein of LHCII type I, chloroplastic* |
| P20507 | *Carbonic anhydrase 1* |
| Q1WLX1 | *Elongation factor Tu, chloroplastic_1* |
| Q5DM57 | *Intraflagellar transport protein 172* |
| Q6DN05 | *Betaine lipid synthase* |
| Q6IYG3 | *NAR1.5* |
| Q6J214 | *15-cis-phytoene synthase* |
| Q6RCE1 | *Intraflagellar transport protein 74* |
| Q6UBQ3 | *Flagellar radial spoke protein 2* |
| Q75VY4 | *Chlorophyll a-b binding protein, chloroplastic_1* |
| Q75VZ0 | *Chlorophyll a-b binding protein, chloroplastic_2* |
| Q8S3T9 | *Chlorophyll a-b binding protein, chloroplastic_3* |
| Q9FDV9 | *4-alpha-glucanotransferase* |
| Q9FYU1 | *Fe-hydrogenase* |
| Q9LD42 | *Fe-assimilating protein 1* |
| Q9SPI9 | *Photosystem II reaction center W protein, chloroplastic* |

| **Table S2: Gene ontology (GO) categories enriched in the differently expressed protein (DEPs) in *Chlamydomonas reinhardti*i strain CC-125 in response to Blue Vs Red light** | | | | | | | | | | | | |
| --- | --- | --- | --- | --- | --- | --- | --- | --- | --- | --- | --- | --- |
| **GO ID** | **GO Name** | **GO Category** | | **Size** | | **Nominal p-val** | | **FDR q-val** | | **FWER p-val** | | **Rank at Max** |
| GO:0007444 | imaginal disc development | BIOLOGICAL_PROCESS | | 15 | | 0.003649635 | | 0.034430217 | | 0.923 | | 1 |
| GO:0045664 | regulation of neuron differentiation | BIOLOGICAL_PROCESS | | 20 | | 0.003669725 | | 0.05081171 | | 0.997 | | 1 |
| GO:0140241 | translation at synapse | BIOLOGICAL_PROCESS | | 20 | | 0.005586592 | | 0.044827897 | | 0.991 | | 2 |
| GO:0043934 | sporulation | BIOLOGICAL_PROCESS | | 19 | | 0 | | 0.01656279 | | 0.437 | | 10 |
| GO:0000470 | maturation of LSU-rRNA | BIOLOGICAL_PROCESS | | 26 | | 0 | | 0.018597325 | | 0.503 | | 22 |
| GO:0000463 | maturation of LSU-rRNA from tricistronic rRNA transcript (SSU-rRNA, 5.8S rRNA, LSU-rRNA) | BIOLOGICAL_PROCESS | | 19 | | 0.003773585 | | 0.024178177 | | 0.738 | | 22 |
| GO:0032984 | protein-containing complex disassembly | BIOLOGICAL_PROCESS | | 36 | | 0 | | 0.018280908 | | 0.518 | | 24 |
| GO:0040025 | vulval development | BIOLOGICAL_PROCESS | | 16 | | 0 | | 0.014611481 | | 0.312 | | 37 |
| GO:0031346 | positive regulation of cell projection organization | BIOLOGICAL_PROCESS | | 36 | | 0 | | 0.023532892 | | 0.709 | | 45 |
| GO:0048592 | eye morphogenesis | BIOLOGICAL_PROCESS | | 23 | | 0 | | 0.04528958 | | 0.99 | | 45 |
| GO:0006400 | tRNA modification | BIOLOGICAL_PROCESS | | 25 | | 0.003649635 | | 0.035547886 | | 0.916 | | 58 |
| GO:0007423 | sensory organ development | BIOLOGICAL_PROCESS | | 66 | | 0.001689189 | | 0.04387467 | | 0.987 | | 153 |
| GO:0002753 | cytoplasmic pattern recognition receptor signaling pathway | BIOLOGICAL_PROCESS | | 15 | | 0.001886793 | | 0.024038985 | | 0.743 | | 167 |
| GO:1902115 | regulation of organelle assembly | BIOLOGICAL_PROCESS | | 25 | | 0.005524862 | | 0.054849047 | | 0.999 | | 256 |
| GO:0002164 | larval development | BIOLOGICAL_PROCESS | | 45 | | 0 | | 0.006998293 | | 0.088 | | 296 |
| GO:0002119 | nematode larval development | BIOLOGICAL_PROCESS | | 33 | | 0 | | 0.014630337 | | 0.305 | | 296 |
| GO:0072331 | signal transduction by p53 class mediator | BIOLOGICAL_PROCESS | | 21 | | 0.001841621 | | 0.03560941 | | 0.915 | | 309 |
| GO:0044089 | positive regulation of cellular component biogenesis | BIOLOGICAL_PROCESS | | 50 | | 0.001760563 | | 0.012968075 | | 0.262 | | 310 |
| GO:0045944 | positive regulation of transcription by RNA polymerase II | BIOLOGICAL_PROCESS | | 49 | | 0 | | 0.0319789 | | 0.869 | | 318 |
| GO:0042391 | regulation of membrane potential | BIOLOGICAL_PROCESS | | 29 | | 0 | | 0.054851875 | | 0.999 | | 327 |
| GO:0051130 | positive regulation of cellular component organization | BIOLOGICAL_PROCESS | | 124 | | 0 | | 0.01581813 | | 0.362 | | 336 |
| GO:0044087 | regulation of cellular component biogenesis | BIOLOGICAL_PROCESS | | 96 | | 0 | | 0.00649605 | | 0.059 | | 348 |
| GO:1902532 | negative regulation of intracellular signal transduction | BIOLOGICAL_PROCESS | | 63 | | 0 | | 0.015262969 | | 0.336 | | 348 |
| GO:0072384 | organelle transport along microtubule | BIOLOGICAL_PROCESS | | 18 | | 0.011111111 | | 0.05222087 | | 0.997 | | 362 |
| GO:0006835 | dicarboxylic acid transport | BIOLOGICAL_PROCESS | | 18 | | 0.003669725 | | 0.050642896 | | 0.997 | | 367 |
| GO:0031401 | positive regulation of protein modification process | BIOLOGICAL_PROCESS | | 40 | | 0 | | 0.015445465 | | 0.367 | | 373 |
| GO:1901699 | cellular response to nitrogen compound | BIOLOGICAL_PROCESS | | 71 | | 0 | | 0.018528214 | | 0.532 | | 373 |
| GO:0030522 | intracellular receptor signaling pathway | BIOLOGICAL_PROCESS | | 34 | | 0 | | 0.029171646 | | 0.815 | | 373 |
| GO:0032103 | positive regulation of response to external stimulus | BIOLOGICAL_PROCESS | | 55 | | 0.001808318 | | 0.031763792 | | 0.869 | | 373 |
| GO:0002221 | pattern recognition receptor signaling pathway | BIOLOGICAL_PROCESS | | 32 | | 0.003496504 | | 0.031835396 | | 0.858 | | 373 |
| GO:0062207 | regulation of pattern recognition receptor signaling pathway | BIOLOGICAL_PROCESS | | 18 | | 0.003676471 | | 0.034943283 | | 0.92 | | 373 |
| GO:0010508 | positive regulation of autophagy | BIOLOGICAL_PROCESS | | 23 | | 0.003703704 | | 0.01889381 | | 0.557 | | 394 |
| GO:0016241 | regulation of macroautophagy | BIOLOGICAL_PROCESS | | 28 | | 0.005376344 | | 0.042574935 | | 0.98 | | 394 |
| GO:0000281 | mitotic cytokinesis | BIOLOGICAL_PROCESS | | 33 | | 0.009140768 | | 0.04945846 | | 0.996 | | 394 |
| GO:0050808 | synapse organization | BIOLOGICAL_PROCESS | | 40 | | 0.003642987 | | 0.05202101 | | 0.997 | | 394 |
| GO:0050727 | regulation of inflammatory response | BIOLOGICAL_PROCESS | | 23 | | 0 | | 0.013801309 | | 0.24 | | 400 |
| GO:0031348 | negative regulation of defense response | BIOLOGICAL_PROCESS | | 28 | | 0.001808318 | | 0.024102436 | | 0.748 | | 400 |
| GO:0036503 | ERAD pathway | BIOLOGICAL_PROCESS | | 18 | | 0.001811594 | | 0.035212375 | | 0.919 | | 400 |
| GO:0050777 | negative regulation of immune response | BIOLOGICAL_PROCESS | | 17 | | 0.009057971 | | 0.045498908 | | 0.989 | | 400 |
| GO:1903825 | organic acid transmembrane transport | BIOLOGICAL_PROCESS | | 33 | | 0.007462686 | | 0.052697137 | | 0.997 | | 402 |
| GO:0031347 | regulation of defense response | BIOLOGICAL_PROCESS | | 91 | | 0 | | 0.001523408 | | 0.001 | | 411 |
| GO:0009895 | negative regulation of catabolic process | BIOLOGICAL_PROCESS | | 48 | | 0.001795332 | | 0.013333689 | | 0.254 | | 411 |
| GO:0051128 | regulation of cellular component organization | BIOLOGICAL_PROCESS | | 255 | | 0 | | 0.052697 | | 0.997 | | 411 |
| GO:1903047 | mitotic cell cycle process | BIOLOGICAL_PROCESS | | 118 | | 0 | | 0.052946307 | | 0.997 | | 427 |
| GO:0042325 | regulation of phosphorylation | BIOLOGICAL_PROCESS | | 47 | | 0 | | 0.007286069 | | 0.087 | | 439 |
| GO:0016310 | phosphorylation | BIOLOGICAL_PROCESS | | 115 | | 0 | | 0.043097284 | | 0.981 | | 439 |
| GO:0080134 | regulation of response to stress | BIOLOGICAL_PROCESS | | 177 | | 0 | | 0.004973938 | | 0.042 | | 442 |
| GO:0006954 | inflammatory response | BIOLOGICAL_PROCESS | | 41 | | 0 | | 0.008992149 | | 0.144 | | 442 |
| GO:0002683 | negative regulation of immune system process | BIOLOGICAL_PROCESS | | 34 | | 0.001908397 | | 0.018074168 | | 0.512 | | 442 |
| GO:0010506 | regulation of autophagy | BIOLOGICAL_PROCESS | | 52 | | 0 | | 0.004744168 | | 0.037 | | 444 |
| GO:0045934 | negative regulation of nucleobase-containing compound metabolic process | BIOLOGICAL_PROCESS | | 110 | | 0.001642036 | | 0.045216776 | | 0.991 | | 466 |
| GO:0048585 | negative regulation of response to stimulus | BIOLOGICAL_PROCESS | | 140 | | 0 | | 0.016626405 | | 0.445 | | 470 |
| GO:0016032 | viral process | BIOLOGICAL_PROCESS | | 36 | | 0 | | 0.04201128 | | 0.976 | | 483 |
| GO:0031344 | regulation of cell projection organization | BIOLOGICAL_PROCESS | | 57 | | 0 | | 0.018167801 | | 0.539 | | 503 |
| GO:0120035 | regulation of plasma membrane bounded cell projection organization | BIOLOGICAL_PROCESS | | 53 | | 0.001782531 | | 0.018512854 | | 0.506 | | 503 |
| GO:0010975 | regulation of neuron projection development | BIOLOGICAL_PROCESS | | 43 | | 0.007104796 | | 0.045273736 | | 0.989 | | 503 |
| GO:0061024 | membrane organization | BIOLOGICAL_PROCESS | | 126 | | 0 | | 0.051796883 | | 0.997 | | 504 |
| GO:0016236 | macroautophagy | BIOLOGICAL_PROCESS | | 62 | | 0 | | 0.01685179 | | 0.462 | | 507 |
| GO:0061919 | process utilizing autophagic mechanism | BIOLOGICAL_PROCESS | | 88 | | 0 | | 0.04480957 | | 0.991 | | 507 |
| GO:0006914 | autophagy | BIOLOGICAL_PROCESS | | 88 | | 0.001808318 | | 0.047295786 | | 0.995 | | 507 |
| GO:0032101 | regulation of response to external stimulus | BIOLOGICAL_PROCESS | | 101 | | 0 | | 0.002291618 | | 0.003 | | 511 |
| GO:0019220 | regulation of phosphate metabolic process | BIOLOGICAL_PROCESS | | 73 | | 0 | | 0.00287056 | | 0.015 | | 511 |
| GO:0051174 | regulation of phosphorus metabolic process | BIOLOGICAL_PROCESS | | 74 | | 0 | | 0.003063822 | | 0.014 | | 511 |
| GO:0002831 | regulation of response to biotic stimulus | BIOLOGICAL_PROCESS | | 76 | | 0 | | 0.006356449 | | 0.066 | | 511 |
| GO:0010562 | positive regulation of phosphorus metabolic process | BIOLOGICAL_PROCESS | | 41 | | 0 | | 0.013213066 | | 0.259 | | 511 |
| GO:0045937 | positive regulation of phosphate metabolic process | BIOLOGICAL_PROCESS | | 41 | | 0 | | 0.013453978 | | 0.242 | | 511 |
| GO:0031399 | regulation of protein modification process | BIOLOGICAL_PROCESS | | 68 | | 0 | | 0.015864417 | | 0.406 | | 511 |
| GO:0002218 | activation of innate immune response | BIOLOGICAL_PROCESS | | 35 | | 0.003780718 | | 0.016008977 | | 0.389 | | 511 |
| GO:0001932 | regulation of protein phosphorylation | BIOLOGICAL_PROCESS | | 39 | | 0.006956522 | | 0.019523997 | | 0.591 | | 511 |
| GO:0051094 | positive regulation of developmental process | BIOLOGICAL_PROCESS | | 105 | | 0 | | 0.029805435 | | 0.83 | | 511 |
| GO:0050776 | regulation of immune response | BIOLOGICAL_PROCESS | | 77 | | 0.00331675 | | 0.03149672 | | 0.858 | | 511 |
| GO:0042327 | positive regulation of phosphorylation | BIOLOGICAL_PROCESS | | 29 | | 0.003454231 | | 0.03532739 | | 0.917 | | 511 |
| GO:0043085 | positive regulation of catalytic activity | BIOLOGICAL_PROCESS | | 38 | | 0.001751314 | | 0.039966032 | | 0.963 | | 511 |
| GO:0001934 | positive regulation of protein phosphorylation | BIOLOGICAL_PROCESS | | 24 | | 0.007117438 | | 0.04003127 | | 0.961 | | 511 |
| GO:0048584 | positive regulation of response to stimulus | BIOLOGICAL_PROCESS | | 186 | | 0 | | 0.040148072 | | 0.959 | | 511 |
| GO:0050865 | regulation of cell activation | BIOLOGICAL_PROCESS | | 18 | | 0.011560693 | | 0.046801724 | | 0.995 | | 511 |
| GO:0002684 | positive regulation of immune system process | BIOLOGICAL_PROCESS | | 74 | | 0 | | 0.05166705 | | 0.997 | | 511 |
| GO:0006955 | immune response | BIOLOGICAL_PROCESS | | 126 | | 0 | | 0.054579094 | | 0.999 | | 511 |
| GO:0009408 | response to heat | BIOLOGICAL_PROCESS | | 86 | | 0 | | 0.039720643 | | 0.953 | | 514 |
| GO:0006909 | phagocytosis | BIOLOGICAL_PROCESS | | 26 | | 0.001730104 | | 0.04204365 | | 0.975 | | 516 |
| GO:0051049 | regulation of transport | BIOLOGICAL_PROCESS | | 171 | | 0 | | 0.019311344 | | 0.593 | | 517 |
| GO:0009723 | response to ethylene | BIOLOGICAL_PROCESS | | 17 | | 0.005524862 | | 0.05404915 | | 0.998 | | 520 |
| GO:0044093 | positive regulation of molecular function | BIOLOGICAL_PROCESS | | 67 | | 0.003649635 | | 0.03978147 | | 0.954 | | 529 |
| GO:0023056 | positive regulation of signaling | BIOLOGICAL_PROCESS | | 149 | | 0 | | 0.04039845 | | 0.968 | | 529 |
| GO:0010647 | positive regulation of cell communication | BIOLOGICAL_PROCESS | | 148 | | 0 | | 0.042946264 | | 0.985 | | 529 |
| GO:1902531 | regulation of intracellular signal transduction | BIOLOGICAL_PROCESS | | 162 | | 0 | | 0.045165222 | | 0.991 | | 529 |
| GO:1902533 | positive regulation of intracellular signal transduction | BIOLOGICAL_PROCESS | | 83 | | 0.005093379 | | 0.05037347 | | 0.997 | | 529 |
| GO:0055088 | lipid homeostasis | BIOLOGICAL_PROCESS | | 25 | | 0.007619048 | | 0.052200686 | | 0.997 | | 533 |
| GO:0002181 | cytoplasmic translation | BIOLOGICAL_PROCESS | | 90 | | 0 | | 0.007618435 | | 0.103 | | 539 |
| GO:0048232 | male gamete generation | BIOLOGICAL_PROCESS | | 97 | | 0 | | 0.04164205 | | 0.974 | | 550 |
| GO:0023057 | negative regulation of signaling | BIOLOGICAL_PROCESS | | 116 | | 0 | | 0.023305329 | | 0.715 | | 555 |
| GO:0009968 | negative regulation of signal transduction | BIOLOGICAL_PROCESS | | 111 | | 0 | | 0.024685834 | | 0.754 | | 555 |
| GO:0010648 | negative regulation of cell communication | BIOLOGICAL_PROCESS | | 116 | | 0 | | 0.029103654 | | 0.816 | | 555 |
| GO:0051402 | neuron apoptotic process | BIOLOGICAL_PROCESS | | 22 | | 0.007380074 | | 0.0516739 | | 0.997 | | 555 |
| GO:0042592 | homeostatic process | BIOLOGICAL_PROCESS | | 208 | | 0 | | 0.00398237 | | 0.013 | | 558 |
| GO:0002682 | regulation of immune system process | BIOLOGICAL_PROCESS | | 111 | | 0 | | 0.016112495 | | 0.424 | | 558 |
| GO:0031349 | positive regulation of defense response | BIOLOGICAL_PROCESS | | 49 | | 0 | | 0.035613578 | | 0.908 | | 558 |
| GO:0015711 | organic anion transport | BIOLOGICAL_PROCESS | | 63 | | 0 | | 0.019380402 | | 0.58 | | 593 |
| GO:0046942 | carboxylic acid transport | BIOLOGICAL_PROCESS | | 46 | | 0.007259528 | | 0.04511254 | | 0.993 | | 593 |
| GO:0015849 | organic acid transport | BIOLOGICAL_PROCESS | | 46 | | 0.003436426 | | 0.045542315 | | 0.989 | | 593 |
| GO:0048878 | chemical homeostasis | BIOLOGICAL_PROCESS | | 136 | | 0 | | 0.007006744 | | 0.08 | | 604 |
| GO:0019725 | cellular homeostasis | BIOLOGICAL_PROCESS | | 120 | | 0 | | 0.014927649 | | 0.336 | | 604 |
| GO:0055082 | intracellular chemical homeostasis | BIOLOGICAL_PROCESS | | 97 | | 0 | | 0.034622703 | | 0.92 | | 604 |
| GO:0006970 | response to osmotic stress | BIOLOGICAL_PROCESS | | 97 | | 0 | | 0.0430865 | | 0.983 | | 605 |
| GO:0032879 | regulation of localization | BIOLOGICAL_PROCESS | | 234 | | 0 | | 0.016600644 | | 0.452 | | 612 |
| GO:0051050 | positive regulation of transport | BIOLOGICAL_PROCESS | | 97 | | 0 | | 0.022552488 | | 0.675 | | 612 |
| GO:0055085 | transmembrane transport | BIOLOGICAL_PROCESS | | 230 | | 0 | | 0.0426782 | | 0.986 | | 612 |
| GO:0032787 | monocarboxylic acid metabolic process | BIOLOGICAL_PROCESS | | 146 | | 0 | | 0.05173799 | | 0.997 | | 615 |
| GO:0034655 | nucleobase-containing compound catabolic process | BIOLOGICAL_PROCESS | | 135 | | 0 | | 0.01819167 | | 0.536 | | 618 |
| GO:0140053 | mitochondrial gene expression | BIOLOGICAL_PROCESS | | 39 | | 0.00177305 | | 0.021257056 | | 0.648 | | 618 |
| GO:0006401 | RNA catabolic process | BIOLOGICAL_PROCESS | | 104 | | 0 | | 0.052331463 | | 0.997 | | 618 |
| GO:0141188 | nucleic acid catabolic process | BIOLOGICAL_PROCESS | | 105 | | 0.003205128 | | 0.052514248 | | 0.997 | | 618 |
| GO:0009894 | regulation of catabolic process | BIOLOGICAL_PROCESS | | 168 | | 0 | | 0.003409589 | | 0.02 | | 631 |
| GO:0043066 | negative regulation of apoptotic process | BIOLOGICAL_PROCESS | | 64 | | 0 | | 0.016295392 | | 0.388 | | 631 |
| GO:0043069 | negative regulation of programmed cell death | BIOLOGICAL_PROCESS | | 73 | | 0 | | 0.019383693 | | 0.599 | | 631 |
| GO:0043067 | regulation of programmed cell death | BIOLOGICAL_PROCESS | | 122 | | 0 | | 0.028565073 | | 0.798 | | 631 |
| GO:0009892 | negative regulation of metabolic process | BIOLOGICAL_PROCESS | | 330 | | 0 | | 0.031518996 | | 0.859 | | 631 |
| GO:0042981 | regulation of apoptotic process | BIOLOGICAL_PROCESS | | 109 | | 0 | | 0.03218143 | | 0.858 | | 631 |
| GO:0032880 | regulation of protein localization | BIOLOGICAL_PROCESS | | 104 | | 0 | | 0.032400656 | | 0.878 | | 631 |
| GO:0080135 | regulation of cellular response to stress | BIOLOGICAL_PROCESS | | 71 | | 0.001712329 | | 0.051550966 | | 0.997 | | 631 |
| GO:0010608 | post-transcriptional regulation of gene expression | BIOLOGICAL_PROCESS | | 165 | | 0 | | 0.016367925 | | 0.423 | | 665 |
| GO:0051247 | positive regulation of protein metabolic process | BIOLOGICAL_PROCESS | | 106 | | 0 | | 0.039445285 | | 0.953 | | 674 |
| GO:0032102 | negative regulation of response to external stimulus | BIOLOGICAL_PROCESS | | 32 | | 0.001834862 | | 0.034377623 | | 0.921 | | 732 |
| GO:0031667 | response to nutrient levels | BIOLOGICAL_PROCESS | | 141 | | 0.001612903 | | 0.045344956 | | 0.991 | | 770 |
| GO:0007417 | central nervous system development | BIOLOGICAL_PROCESS | | 86 | | 0 | | 0.045408186 | | 0.989 | | 818 |
| GO:0006913 | nucleocytoplasmic transport | BIOLOGICAL_PROCESS | | 78 | | 0.003401361 | | 0.035176165 | | 0.927 | | 828 |
| GO:0051169 | nuclear transport | BIOLOGICAL_PROCESS | | 78 | | 0.001618123 | | 0.03999125 | | 0.96 | | 828 |
| GO:0071705 | nitrogen compound transport | BIOLOGICAL_PROCESS | | 299 | | 0 | | 0.012735688 | | 0.264 | | 835 |
| GO:0051246 | regulation of protein metabolic process | BIOLOGICAL_PROCESS | | 215 | | 0 | | 0.006254837 | | 0.068 | | 851 |
| GO:0006417 | regulation of translation | BIOLOGICAL_PROCESS | | 103 | | 0 | | 0.015699126 | | 0.364 | | 868 |
| GO:0051248 | negative regulation of protein metabolic process | BIOLOGICAL_PROCESS | | 70 | | 0.003448276 | | 0.019302487 | | 0.571 | | 868 |
| GO:0006412 | translation | BIOLOGICAL_PROCESS | | 265 | | 0 | | 0.006268135 | | 0.061 | | 879 |
| GO:0005576 | extracellular region | CELLULAR_COMPONENT | | 94 | | 0 | | 0.003574459 | | 0.014 | | 754 |
| GO:0022626 | cytosolic ribosome | CELLULAR_COMPONENT | | 72 | | 0 | | 0.003831639 | | 0.025 | | 498 |
| GO:0022625 | cytosolic large ribosomal subunit | CELLULAR_COMPONENT | | 34 | | 0 | | 0.004590499 | | 0.009 | | 48 |
| GO:0044391 | ribosomal subunit | CELLULAR_COMPONENT | | 72 | | 0 | | 0.004757937 | | 0.034 | | 498 |
| GO:0005840 | ribosome | CELLULAR_COMPONENT | | 107 | | 0 | | 0.007619377 | | 0.099 | | 498 |
| GO:0015934 | large ribosomal subunit | CELLULAR_COMPONENT | | 40 | | 0 | | 0.008180616 | | 0.125 | | 411 |
| GO:0098588 | bounding membrane of organelle | CELLULAR_COMPONENT | | 330 | | 0 | | 0.014236827 | | 0.312 | | 563 |
| GO:0045202 | synapse | CELLULAR_COMPONENT | | 225 | | 0 | | 0.015500127 | | 0.397 | | 507 |
| GO:0098590 | plasma membrane region | CELLULAR_COMPONENT | | 99 | | 0 | | 0.01570136 | | 0.389 | | 544 |
| GO:0030141 | secretory granule | CELLULAR_COMPONENT | | 75 | | 0 | | 0.018480426 | | 0.536 | | 677 |
| GO:0030054 | cell junction | CELLULAR_COMPONENT | | 247 | | 0 | | 0.02229322 | | 0.675 | | 563 |
| GO:0005615 | extracellular space | CELLULAR_COMPONENT | | 37 | | 0.003603604 | | 0.024105458 | | 0.74 | | 448 |
| GO:0005789 | endoplasmic reticulum membrane | CELLULAR_COMPONENT | | 106 | | 0 | | 0.029028818 | | 0.808 | | 716 |
| GO:0098827 | endoplasmic reticulum subcompartment | CELLULAR_COMPONENT | | 107 | | 0.001669449 | | 0.032292604 | | 0.869 | | 716 |
| GO:0099503 | secretory vesicle | CELLULAR_COMPONENT | | 96 | | 0 | | 0.035975534 | | 0.911 | | 612 |
| GO:0005886 | plasma membrane | CELLULAR_COMPONENT | | 316 | | 0 | | 0.037316725 | | 0.941 | | 520 |
| GO:0030139 | endocytic vesicle | CELLULAR_COMPONENT | | 45 | | 0.001848429 | | 0.03994061 | | 0.963 | | 771 |
| GO:0016469 | proton-transporting two-sector ATPase complex | CELLULAR_COMPONENT | | 22 | | 0.005524862 | | 0.040454835 | | 0.97 | | 296 |
| GO:0035770 | ribonucleoprotein granule | CELLULAR_COMPONENT | | 85 | | 0 | | 0.043108113 | | 0.983 | | 693 |
| GO:0042579 | microbody | CELLULAR_COMPONENT | | 58 | | 0.001808318 | | 0.04699485 | | 0.995 | | 791 |
| GO:0099080 | supramolecular complex | CELLULAR_COMPONENT | | 184 | | 0 | | 0.04724027 | | 0.995 | | 416 |
| GO:0042175 | nuclear outer membrane-endoplasmic reticulum membrane network | CELLULAR_COMPONENT | | 112 | | 0 | | 0.04897313 | | 0.996 | | 716 |
| GO:0005777 | peroxisome | CELLULAR_COMPONENT | | 58 | | 0.001647446 | | 0.05221959 | | 0.997 | | 791 |
| GO:0003729 | mRNA binding | MOLECULAR_FUNCTION | | 202 | | 0 | | 0.003838545 | | 0.01 | | 868 |
| GO:0005215 | transporter activity | MOLECULAR_FUNCTION | | 194 | | 0 | | 0.007553526 | | 0.105 | | 604 |
| GO:0022804 | active transmembrane transporter activity | MOLECULAR_FUNCTION | | 113 | | 0 | | 0.007816047 | | 0.114 | | 604 |
| GO:0022857 | transmembrane transporter activity | MOLECULAR_FUNCTION | | 177 | | 0 | | 0.009845628 | | 0.161 | | 604 |
| GO:0022890 | inorganic cation transmembrane transporter activity | MOLECULAR_FUNCTION | | 72 | | 0 | | 0.010656543 | | 0.179 | | 410 |
| GO:0015318 | inorganic molecular entity transmembrane transporter activity | MOLECULAR_FUNCTION | | 96 | | 0 | | 0.012251748 | | 0.209 | | 597 |
| GO:0008514 | organic anion transmembrane transporter activity | MOLECULAR_FUNCTION | | 59 | | 0 | | 0.013702673 | | 0.253 | | 612 |
| GO:0015078 | proton transmembrane transporter activity | MOLECULAR_FUNCTION | | 45 | | 0 | | 0.01550085 | | 0.391 | | 296 |
| GO:0045182 | translation regulator activity | MOLECULAR_FUNCTION | | 63 | | 0.001727116 | | 0.018329196 | | 0.51 | | 870 |
| GO:0022853 | active monoatomic ion transmembrane transporter activity | MOLECULAR_FUNCTION | | 51 | | 0 | | 0.021530312 | | 0.648 | | 296 |
| GO:0030234 | enzyme regulator activity | MOLECULAR_FUNCTION | | 119 | | 0 | | 0.02220158 | | 0.68 | | 672 |
| GO:0090079 | translation regulator activity, nucleic acid binding | MOLECULAR_FUNCTION | | 55 | | 0 | | 0.023285428 | | 0.721 | | 665 |
| GO:0042626 | ATPase-coupled transmembrane transporter activity | MOLECULAR_FUNCTION | | 55 | | 0.001689189 | | 0.023525773 | | 0.713 | | 612 |
| GO:0008324 | monoatomic cation transmembrane transporter activity | MOLECULAR_FUNCTION | | 77 | | 0 | | 0.030018874 | | 0.833 | | 444 |
| GO:0015291 | secondary active transmembrane transporter activity | MOLECULAR_FUNCTION | | 45 | | 0.001818182 | | 0.03238783 | | 0.867 | | 597 |
| GO:0008135 | translation factor activity, RNA binding | MOLECULAR_FUNCTION | | 50 | | 0.005208334 | | 0.035656337 | | 0.911 | | 665 |
| GO:0016810 | hydrolase activity, acting on carbon-nitrogen (but not peptide) bonds | MOLECULAR_FUNCTION | | 32 | | 0.005424955 | | 0.037538763 | | 0.941 | | 527 |
| GO:0015075 | monoatomic ion transmembrane transporter activity | MOLECULAR_FUNCTION | | 92 | | 0 | | 0.040057626 | | 0.962 | | 597 |
| GO:0042802 | identical protein binding | MOLECULAR_FUNCTION | | 284 | | 0 | | 0.040149033 | | 0.964 | | 699 |
| GO:0043021 | ribonucleoprotein complex binding | MOLECULAR_FUNCTION | | 75 | | 0.001584786 | | 0.040154807 | | 0.963 | | 673 |
| GO:0015399 | primary active transmembrane transporter activity | MOLECULAR_FUNCTION | | 67 | | 0.001655629 | | 0.040499978 | | 0.967 | | 604 |
| GO:0060090 | molecular adaptor activity | MOLECULAR_FUNCTION | | 113 | | 0 | | 0.041734457 | | 0.975 | | 427 |
| GO:0019829 | ATPase-coupled monoatomic cation transmembrane transporter activity | MOLECULAR_FUNCTION | | 25 | | 0.007407407 | | 0.041777007 | | 0.976 | | 544 |
| GO:0030695 | GTPase regulator activity | MOLECULAR_FUNCTION | | 28 | | 0.001872659 | | 0.042095035 | | 0.976 | | 503 |
| GO:0016874 | ligase activity | MOLECULAR_FUNCTION | | 91 | | 0 | | 0.04281067 | | 0.979 | | 894 |
| GO:0046943 | carboxylic acid transmembrane transporter activity | MOLECULAR_FUNCTION | | 39 | | 0.00528169 | | 0.042844843 | | 0.983 | | 612 |
| GO:0005342 | organic acid transmembrane transporter activity | MOLECULAR_FUNCTION | | 39 | | 0.003436426 | | 0.04292422 | | 0.983 | | 612 |
| GO:0098772 | molecular function regulator activity | MOLECULAR_FUNCTION | | 166 | | 0 | | 0.04505469 | | 0.991 | | 672 |
| GO:0003735 | structural constituent of ribosome | MOLECULAR_FUNCTION | | 82 | | 0.003372681 | | 0.047165245 | | 0.995 | | 498 |
| GO:0005198 | structural molecule activity | MOLECULAR_FUNCTION | | 124 | | 0.00166113 | | 0.05183077 | | 0.997 | | 417 |

**Table S3. List of important nodes present in bio-curated PPI with their functions as obtained from String database.**

| **Nodes** | ***Functions*** |
| --- | --- |
| ***Klebsormidium nitens*** | |
| COP3 | *Uncharacterized protein. (719 aa)* |
| A0A2K3CQK5 | *Uncharacterized protein. (596 aa)* |
| SQD2 | *Uncharacterized protein. (553 aa)* |
| PSY | *Chloroplast phytoene synthase. (382 aa)* |
| LHCSR1 | *Light-harvesting complex stress-related protein 1, chloroplastic; Required for non-photochemical quenching (NPQ), a mechanism that converts and dissipates the harmful excess absorbed light energy into heat and protect the photosynthetic apparatus from photo-oxidative damage. Is able to sense luminal acidification of the thylakoid membranes, which occurs along with elevated electron flow caused by excess light, and to induce a large, fast, and reversible pH-dependent quenching in LHCII- containing membranes. Mediates excitation energy transfer from light-harvesting complex II (LHCII) to [...] (253 aa)* |
| GSA | *Glutamate-1-semialdehyde aminotransferase; Belongs to the class-III pyridoxal-phosphate-dependent aminotransferase family. (463 aa)* |
| A0A2K3E782 | *Chloride channel protein. (969 aa)* |
| UVR3 | *Cryptochrome photoreceptor. (595 aa)* |
| KAS2 | *3-oxoacyl-[acyl-carrier-protein] synthase; Belongs to the thiolase-like superfamily. Beta-ketoacyl-ACP synthases family. (459 aa)* |
| CPH1 | *Cryptochrome photoreceptor. (1008 aa)* |
| LHCSR3.1 | *Light-harvesting complex stress-related protein 3.1, chloroplastic; Required for non-photochemical quenching (NPQ), a mechanism that converts and dissipates the harmful excess absorbed light energy into heat and protect the photosynthetic apparatus from photo-oxidative damage. NPQ includes dissipating excess light energy to heat (qE) and the reversible coupling of LHCII to photosystems (state transitions or qT), which are considered separate NPQ mechanisms. Is responsible for most of the excess light energy to heat dissipation (qE), also known as energy-dependent chlorophyll fluorescen [...] (259 aa)* |
| ZDS1 | *Zeta-carotene desaturase; Catalyzes the conversion of zeta-carotene to lycopene via the intermediary of neurosporene. It carries out two consecutive desaturations (introduction of double bonds) at positions C-7 and C-7'. (582 aa)* |
| rbcL | *Ribulose bisphosphate carboxylase large chain; RuBisCO catalyzes two reactions: the carboxylation of D- ribulose 1,5-bisphosphate, the primary event in carbon dioxide fixation, as well as the oxidative fragmentation of the pentose substrate in the photorespiration process. Both reactions occur simultaneously and in competition at the same active site. (475 aa)* |
| A0A2K3E785 | *Uncharacterized protein. (751 aa)* |
| cop4 | *Chlamyopsin 4 light-gated ion channel. (737 aa)* |
| psaA | *Photosystem I P700 chlorophyll a apoprotein A1; PsaA and PsaB bind P700, the primary electron donor of photosystem I (PSI), as well as the electron acceptors A0, A1 and FX. PSI is a plastocyanin/cytochrome c6-ferredoxin oxidoreductase, converting photonic excitation into a charge separation, which transfers an electron from the donor P700 chlorophyll pair to the spectroscopically characterized acceptors A0, A1, FX, FA and FB in turn. Oxidized P700 is reduced on the lumenal side of the thylakoid membrane by plastocyanin or cytochrome c6. (751 aa)* |
| PDS | *Chloroplast phytoene desaturase. (564 aa)* |
| EIF3G | *Eukaryotic translation initiation factor 3 subunit G; RNA-binding component of the eukaryotic translation initiation factor 3 (eIF-3) complex, which is involved in protein synthesis of a specialized repertoire of mRNAs and, together with other initiation factors, stimulates binding of mRNA and methionyl-tRNAi to the 40S ribosome. The eIF-3 complex specifically targets and initiates translation of a subset of mRNAs involved in cell proliferation. This subunit can bind 18S rRNA. (283 aa)* |
| A0A2K3E780 | *RRM domain-containing protein. (271 aa)* |
| PRP19 | *Spliceosome component, nuclear pre-mRNA splicing factor. (503 aa)* |
| LCYE | *Lycopene epsilon cyclase. (583 aa)* |
| psaB | *Photosystem I P700 chlorophyll a apoprotein A2; PsaA and PsaB bind P700, the primary electron donor of photosystem I (PSI), as well as the electron acceptors A0, A1 and FX. PSI is a plastocyanin/cytochrome c6-ferredoxin oxidoreductase, converting photonic excitation into a charge separation, which transfers an electron from the donor P700 chlorophyll pair to the spectroscopically characterized acceptors A0, A1, FX, FA and FB in turn. Oxidized P700 is reduced on the lumenal side of the thylakoid membrane by plastocyanin or cytochrome c6; Belongs to the PsaA/PsaB family. (735 aa)* |
| PHOT | *Phototropin; Protein kinase that acts as a blue light photoreceptor. Required for non-photochemical quenching (NPQ), a mechanism that converts and dissipates the harmful excess absorbed light energy into heat and protect the photosynthetic apparatus from photo-oxidative damage. Controls the energy-dependent chlorophyll fluorescence quenching (qE) activity of chlorophyll excited states by inducing the expression of the qE effector protein LHCSR3 in high light intensities. (749 aa)* |
| FAT1 | *Acyl-[acyl-carrier-protein] hydrolase; Plays an essential role in chain termination during de novo fatty acid synthesis; Belongs to the acyl-ACP thioesterase family. (395 aa)* |
| ERF1 | *Eukaryotic release factor 1. (438 aa)* |
| A0A2K3D1J8 | *Uncharacterized protein. (734 aa)* |
| FAD7 | *Chloroplast glycerolipid omega-3-fatty acid desaturase. (418 aa)* |
| ***Arabidopsis thaliana*** | |
| F24B9.25 | Histone H4; Core component of nucleosome. Nucleosomes wrap and compact DNA into chromatin, limiting DNA accessibility to the cellular machineries which require DNA as a template. Histones thereby play a central role in transcription regulation, DNA repair, DNA replication and chromosomal stability. DNA accessibility is regulated via a complex set of post-translational modifications of histones, also called histone code, and nucleosome remodeling. (103 aa) |
| ELF6 | Probable lysine-specific demethylase ELF6; Acts probably as a histone 'Lys-4' (H3K4me) demethylase. Involved in transcriptional gene regulation. Acts as a repressor of the photoperiodic flowering pathway and of FT. Binds around the transcription start site of the FT locus. (1340 aa) |
| MBK5.18 | Zinc ion binding / DNA binding protein. (602 aa) |
| NRPB11 | DNA-directed RNA polymerases II, IV and V subunit 11; DNA-dependent RNA polymerase catalyzes the transcription of DNA into RNA using the four ribonucleoside triphosphates as substrates. Component of RNA polymerase II which synthesizes mRNA precursors and many functional non-coding RNAs. Pol II is the central component of the basal RNA polymerase II transcription machinery. It is composed of mobile elements that move relative to each other. NRPB11 is part of the core element with the central large cleft. Component of RNA polymerases IV and V which mediate short-interfering RNAs (siRNA) [...] (116 aa) |
| NRPB4 | DNA-directed RNA polymerase II subunit 4; DNA-dependent RNA polymerase catalyzes the transcription of DNA into RNA using the four ribonucleoside triphosphates as substrates. Second largest component of RNA polymerase II which synthesizes mRNA precursors and many functional non-coding RNAs. Proposed to contribute to the polymerase catalytic activity and forms the polymerase active center together with the largest subunit. Pol II is the central component of the basal RNA polymerase II transcription machinery. It is composed of mobile elements that move relative to each other. (138 aa) |
| PH1 | Pleckstrin homology domain-containing protein 1; Binds specifically to phosphatidylinositol 3-phosphate (PtdIns3P), but not to other phosphoinositides. (145 aa) |
| RBL-2 | Protein RBL; Promotes the expression of FLC and FLC homologs to repress the floral transition. Promotes WRKY70 and LTP7 genes epigenetic methylation (e.g. H3K4me3) and subsequent expression. (547 aa) |
| RPL40B | Ubiquitin-60S ribosomal protein L40-2; [Ubiquitin]: exists either covalently attached to another protein, or free (unanchored). When covalently bound, it is conjugated to target proteins via an isopeptide bond either as a monomer (monoubiquitin), a polymer linked via different Lys residues of the ubiquitin (polyubiquitin chains) or a linear polymer linked via the initiator Met of the ubiquitin (linear polyubiquitin chains). Polyubiquitin chains, when attached to a target protein, have different functions depending on the Lys residue of the ubiquitin that is linked: Lys-11-linked is inv [...] (128 aa) |
| T27K22.4 | annotation not available (824 aa) |
| TRO | Protein TRAUCO; Trithorax-group gene homolog required for early embryogenesis. Required for the expression of FLC and FLC homologs and represses flowering. Required for proper leaf growth and development. Part of COMPASS-like complexes responsible for H3K4 trimethylation, but not for di- or mono-methylation of histone H3 'Lys-4'. Binds to target loci chromatin, increasing H3K4 trimethylation and causing activation of the gene. Involved in the transition from transcription initiation to transcription elongation. (509 aa) |
| CTF2B | FAD/NAD(P)-binding oxidoreductase family protein. (427 aa) |
| PHOT1 | Phototropin-1; Protein kinase that acts as a blue light photoreceptor in a signal-transduction pathway for photo-induced movements. Phosphorylates BLUS1, a kinase involved in stomatal opening. Required for blue light mediated mRNA destabilization. Mediates calcium spiking of extracellular origin in response to a low rate of blue light. Also mediates rapid membrane depolarization and growth inhibition in response to blue light. Necessary for root phototropism. Involved in hypocotyl phototropism under a low rate but not under a high rate of blue light. Contributes to the chloroplast accu [...] (996 aa) |
| PHOT2 | Phototropin-2; Protein kinase that acts as a blue light photoreceptor in a signal-transduction pathway for photo-induced movements. Phosphorylates BLUS1, a kinase involved in stomatal opening. Mediates calcium spiking of extra- and intracellular origins in response to blue light. Involved in hypocotyl phototropism. Contributes to the chloroplast accumulation in low blue light and mediates their translocation (avoidance response) at high fluence. Regulates stomata opening and photomorphogenesis response of leaf tissue. Not involved in hypocotyl elongation inhibition, anthocyanin accumul [...] (915 aa) |
| CRY2 | Cryptochrome-2; Photoreceptor that mediates primarily blue light inhibition of hypocotyl elongation and photoperiodic control of floral initiation, and regulates other light responses, including circadian rhythms, tropic growth, stomata opening, guard cell development, root development, bacterial and viral pathogen responses, abiotic stress responses, cell cycles, programmed cell death, apical dominance, fruit and ovule development, seed dormancy, and magnetoreception. Photoexcited cryptochromes interact with signaling partner proteins to alter gene expression at both transcriptional a [...] (612 aa) |
| CRYD | Cryptochrome DASH, chloroplastic/mitochondrial; May have a photoreceptor function. Binds ss- and ds-DNA in a sequence non-specific manner. Has a photolyase activity specific for cyclobutane pyrimidine dimers in ssDNA; Belongs to the DNA photolyase class-1 family. (569 aa) |
| UVR3 | (6-4)DNA photolyase; Involved in repair of UV radiation-induced DNA damage. Catalyzes the photoreactivation of pyrimidine [6-4] pyrimidone photoproduct (6-4 products). Binds specifically to DNA containing 6-4 products and repairs these lesions in a visible light-dependent manner. Not required for repair of cyclobutane pyrimidine dimer (CPD). (556 aa) |
| PHYE | Phytochrome E; Regulatory photoreceptor which exists in two forms that are reversibly interconvertible by light: the Pr form that absorbs maximally in the red region of the spectrum and the Pfr form that absorbs maximally in the far-red region. Photoconversion of Pr to Pfr induces an array of morphogenic responses, whereas reconversion of Pfr to Pr cancels the induction of those responses. Pfr controls the expression of a number of nuclear genes including those encoding the small subunit of ribulose-bisphosphate carboxylase, chlorophyll A/B binding protein, protochlorophyllide reductas [...] (1112 aa) |
| PHYD | Phytochrome D; Regulatory photoreceptor which exists in two forms that are reversibly interconvertible by light: the Pr form that absorbs maximally in the red region of the spectrum and the Pfr form that absorbs maximally in the far-red region. Photoconversion of Pr to Pfr induces an array of morphogenic responses, whereas reconversion of Pfr to Pr cancels the induction of those responses. Pfr controls the expression of a number of nuclear genes including those encoding the small subunit of ribulose-bisphosphate carboxylase, chlorophyll A/B binding protein, protochlorophyllide reductas [...] (1164 aa) |
| PHYA | Phytochrome A; Regulatory photoreceptor which exists in two forms that are reversibly interconvertible by light: the Pr form that absorbs maximally in the red region of the spectrum and the Pfr form that absorbs maximally in the far-red region. Photoconversion of Pr to Pfr induces an array of morphogenetic responses, whereas reconversion of Pfr to Pr cancels the induction of those responses. Pfr controls the expression of a number of nuclear genes including those encoding the small subunit of ribulose-bisphosphate carboxylase, chlorophyll A/B binding protein, protochlorophyllide reduct [...] (1122 aa) |
| PDS | 15-cis-phytoene desaturase, chloroplastic/chromoplastic; Converts phytoene into zeta-carotene via the intermediary of phytofluene by the symmetrical introduction of two double bonds at the C-11 and C-11' positions of phytoene with a concomitant isomerization of two neighboring double bonds at the C9 and C9' positions from trans to cis; Belongs to the carotenoid/retinoid oxidoreductase family. (566 aa) |
| PSY1 | Phytoene synthase, chloroplastic; Catalyzes the reaction from prephytoene diphosphate to phytoene; Belongs to the phytoene/squalene synthase family. (422 aa) |
| LCY1 | Lycopene beta cyclase, chloroplastic; Involved in carotenoid biosynthesis. Catalyzes the double cyclization reaction which converts lycopene to beta-carotene and neurosporene to beta-zeacarotene. Major lycopene beta- cyclase that does not seem to be involved in neoxanthin synthesis. Involved in salt tolerance improvement by increasing synthesis of carotenoids, which impairs reactive oxygen species (ROS) and protects the photosynthetic system under salt stress. (501 aa) |
| BETA-OHASE_1 | Beta-carotene 3-hydroxylase 1, chloroplastic; Nonheme diiron monooxygenase involved in the biosynthesis of xanthophylls. Specific for beta-ring hydroxylations of beta-carotene. Has also a low activity toward the beta- and epsilon-rings of alpha- carotene. No activity with acyclic carotenoids such as lycopene and neurosporene. Uses ferredoxin as an electron donor. Belongs to the sterol desaturase family. (310 aa) |
| F7D8.18 | Violaxanthin de-epoxidase-like protein. (522 aa) |
| ZEP | Zeaxanthin epoxidase, chloroplastic; Zeaxanthin epoxidase that plays an important role in the xanthophyll cycle and abscisic acid (ABA) biosynthesis. Converts zeaxanthin into antheraxanthin and subsequently violaxanthin. Required for resistance to osmotic and drought stresses, ABA-dependent stomatal closure, seed development and dormancy, modulation of defense gene expression and disease resistance and non-photochemical quencing (NPQ). Through its role in ABA biosynthesis, regulates the expression of stress-responsive genes such as RD29A during osmotic stress and is required for normal [...] (667 aa |
| ***Oryza sativa*** | |
| OsI_17352 | EGF_CA domain-containing protein. (663 aa) |
| OsI_17350 | Uncharacterized protein. (437 aa) |
| OsI_17354 | Uncharacterized protein; Belongs to the terpene synthase family. (697 aa) |
| OsI_17355 | Terpene_synth domain-containing protein. (407 aa) |
| OsI_17356 | Uncharacterized protein; Belongs to the terpene synthase family. (1130 aa) |
| OsI_17357 | Uncharacterized protein; Belongs to the terpene synthase family. (740 aa) |
| OsI_17358 | X8 domain-containing protein. (67 aa) |
| OsI_24748 | Uncharacterized protein. (148 aa) |
| OsI_11521 | Photolyase/cryptochrome alpha/beta domain-containing protein. (451 aa) |
| PHYC | Phytochrome C; Regulatory photoreceptor which exists in two forms that are reversibly interconvertible by light: the Pr form that absorbs maximally in the red region of the spectrum and the Pfr form that absorbs maximally in the far-red region. Photoconversion of Pr to Pfr induces an array of morphogenic responses, whereas reconversion of Pfr to Pr cancels the induction of those responses. Pfr controls the expression of a number of nuclear genes including those encoding the small subunit of ribulose-bisphosphate carboxylase, chlorophyll A/B binding protein, protochlorophyllide reductas [...] (1137 aa) |
| PHYB | Phytochrome B; Regulatory photoreceptor which exists in two forms that are reversibly interconvertible by light: the Pr form that absorbs maximally in the red region of the spectrum and the Pfr form that absorbs maximally in the far-red region. Photoconversion of Pr to Pfr induces an array of morphogenic responses, whereas reconversion of Pfr to Pr cancels the induction of those responses. Pfr controls the expression of a number of nuclear genes including those encoding the small subunit of ribulose-bisphosphate carboxylase, chlorophyll A/B binding protein, protochlorophyllide reductas [...] (1171 aa) |
| OsI_19661 | Uncharacterized protein. (238 aa) |
| PHYA | Phytochrome A; Regulatory photoreceptor which exists in two forms that are reversibly interconvertible by light: the Pr form that absorbs maximally in the red region of the spectrum and the Pfr form that absorbs maximally in the far-red region. Photoconversion of Pr to Pfr induces an array of morphogenic responses, whereas reconversion of Pfr to Pr cancels the induction of those responses. Pfr controls the expression of a number of nuclear genes including those encoding the small subunit of ribulose-bisphosphate carboxylase, chlorophyll A/B binding protein, protochlorophyllide reductas [...] (1128 aa) |
| PDS1 | 15-cis-phytoene desaturase, chloroplastic/chromoplastic; Converts phytoene into zeta-carotene via the intermediary of phytofluene by the symmetrical introduction of two double bonds at the C-11 and C-11' positions of phytoene with a concomitant isomerization of two neighboring double bonds at the C9 and C9' positions from trans to cis. Active with decylplastoquinone (DPQ) as substrate. Also active with other benzoquinones, which are strongly preferred over naphthoquinones as substrates ; Belongs to the carotenoid/retinoid oxidoreductase family. (578 aa) |
| OsI_39199 | Uncharacterized protein. (398 aa) |
| OsI_06183 | Uncharacterized protein. (372 aa) |
| OsI_16086 | Zeaxanthin epoxidase, chloroplastic; Converts zeaxanthin into antheraxanthin and subsequently violaxanthin. (644 aa) |
| OsI_32864 | Uncharacterized protein. (289 aa) |
| OsI_00411 | Uncharacterized protein. (347 aa) |
| OsI_03423 | Uncharacterized protein; Belongs to the FPP/GGPP synthase family. (392 aa) |
| OsI_06748 | Uncharacterized protein; Belongs to the terpene synthase family. (864 aa) |
| OsI_20844 | Uncharacterized protein; Belongs to the FPP/GGPP synthase family. (355 aa) |
| OsI_23427 | Uncharacterized protein; Belongs to the cytochrome P450 family. (486 aa) |
| OsI_23428 | Uncharacterized protein. (69 aa) |
| OsI_23430 | Uncharacterized protein; Belongs to the cytochrome P450 family. (506 aa) |
| OsI_26813 | Alkyl transferase; Belongs to the UPP synthase family. (299 aa) |
| OsI_28830 | Uncharacterized protein. (1206 aa) |
| OsI_30093 | Uncharacterized protein. (709 aa) |
| OsI_30696 | Mediator of RNA polymerase II transcription subunit 14; Component of the Mediator complex, a coactivator involved in the regulated transcription of nearly all RNA polymerase II-dependent genes. Mediator functions as a bridge to convey information from gene- specific regulatory proteins to the basal RNA polymerase II transcription machinery. Mediator is recruited to promoters by direct interactions with regulatory proteins and serves as a scaffold for the assembly of a functional preinitiation complex with RNA polymerase II and the general transcription factors. (1740 aa) |
| OsI_00663 | Uncharacterized protein. (1176 aa) |
| OsI_06757 | Uncharacterized protein. (1086 aa) |
| OsI_14451 | Uncharacterized protein. (245 aa) |
| OsI_14461 | Peptidase A1 domain-containing protein. (193 aa) |
| OsI_23866 | Uncharacterized protein. (265 aa) |
| OsI_26549 | Uncharacterized protein. (522 aa) |
| OsI_27489 | ACT domain-containing protein. (180 aa) |
| OsI_29626 | 14_3_3 domain-containing protein; Belongs to the 14-3-3 family. (264 aa) |
| OsI_32899 | Uncharacterized protein. (450 aa) |
| OsI_36621 | Uncharacterized protein. (1133 aa) |
| OsI_09099 | Nitrate reductase; Nitrate reductase is a key enzyme involved in the first step of nitrate assimilation in plants, fungi and bacteria. (893 aa) |
| OsI_26153 | Thioredoxin domain-containing protein. (432 aa) |
| OsI_28845 | Uncharacterized protein. (1170 aa) |
| OsI_29535 | FAD-binding FR-type domain-containing protein. (434 aa) |
| OsI_29538 | NADH-cytochrome b5 reductase; Belongs to the flavoprotein pyridine nucleotide cytochrome reductase family. (210 aa) |
| OsI_29539 | Uncharacterized protein. (526 aa) |
| OsI_29540 | Cytochrome b5 heme-binding domain-containing protein. (794 aa) |
| OsI_29541 | Uncharacterized protein. (688 aa) |
| OsI_33072 | FAD-binding FR-type domain-containing protein. (547 aa) |
| OsI_36312 | NAD_binding_1 domain-containing protein. (81 aa) |

**Supplementary Figures**

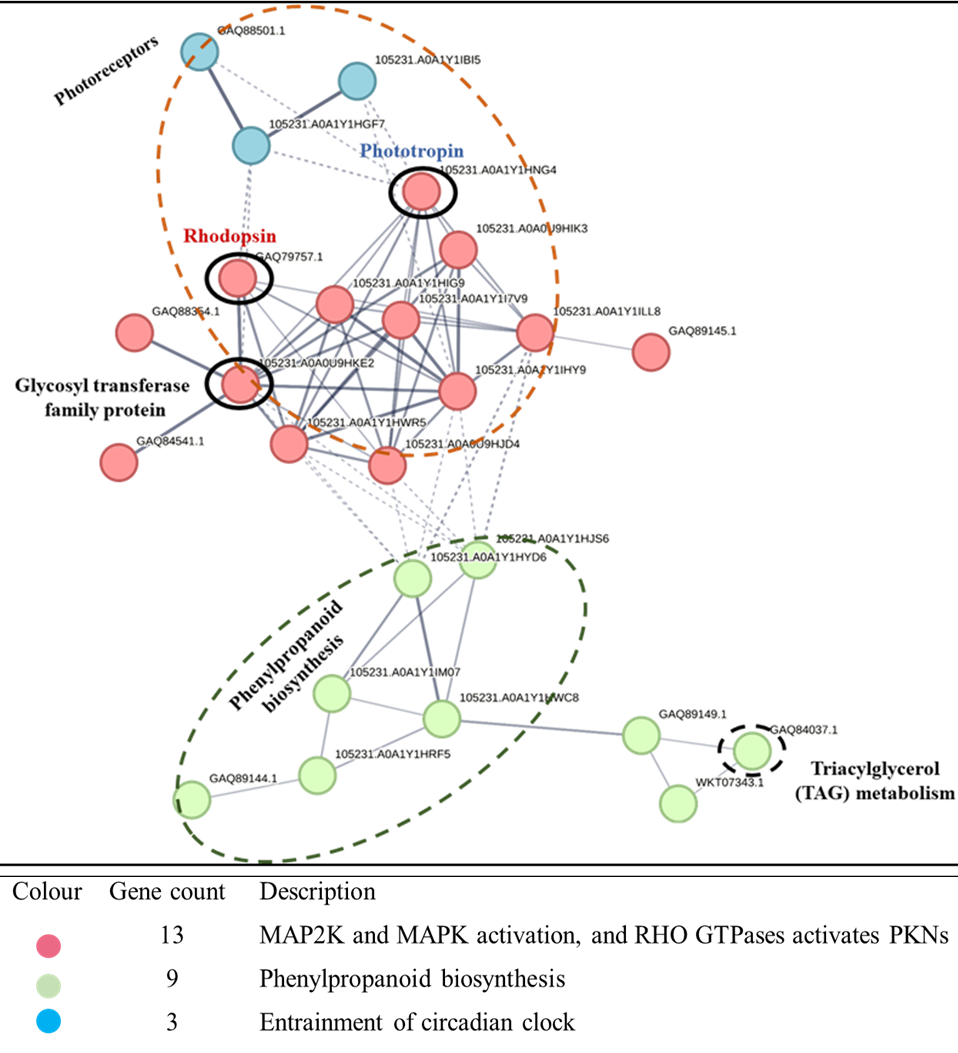
**Supplementary Figure 1**

**Figure S1: Bio-curation of opto-modulation of BGCs for targeted metabolite production in *Klebsormidium nitens***

It shows the bio-curated crosstalk of BGCs cluster and photoreceptor via predicted curated PPI networking in *Klebsormidium nitens*. Here, glycosyl transferase family proteins (BGC core domain component) showed direct interaction with phototropin (a blue light-sensing photoreceptor) and rhodopsin (chlamyopsin), a UV-IR light-sensing photoreceptor (marked in a small dark black colour circle). The k-means clustering method was applied to cluster the PPI. Also, secondary interaction of photoreceptors was observed with phenylpropanoid biosynthesis BGCs component.

**Supplementary Figure 2**

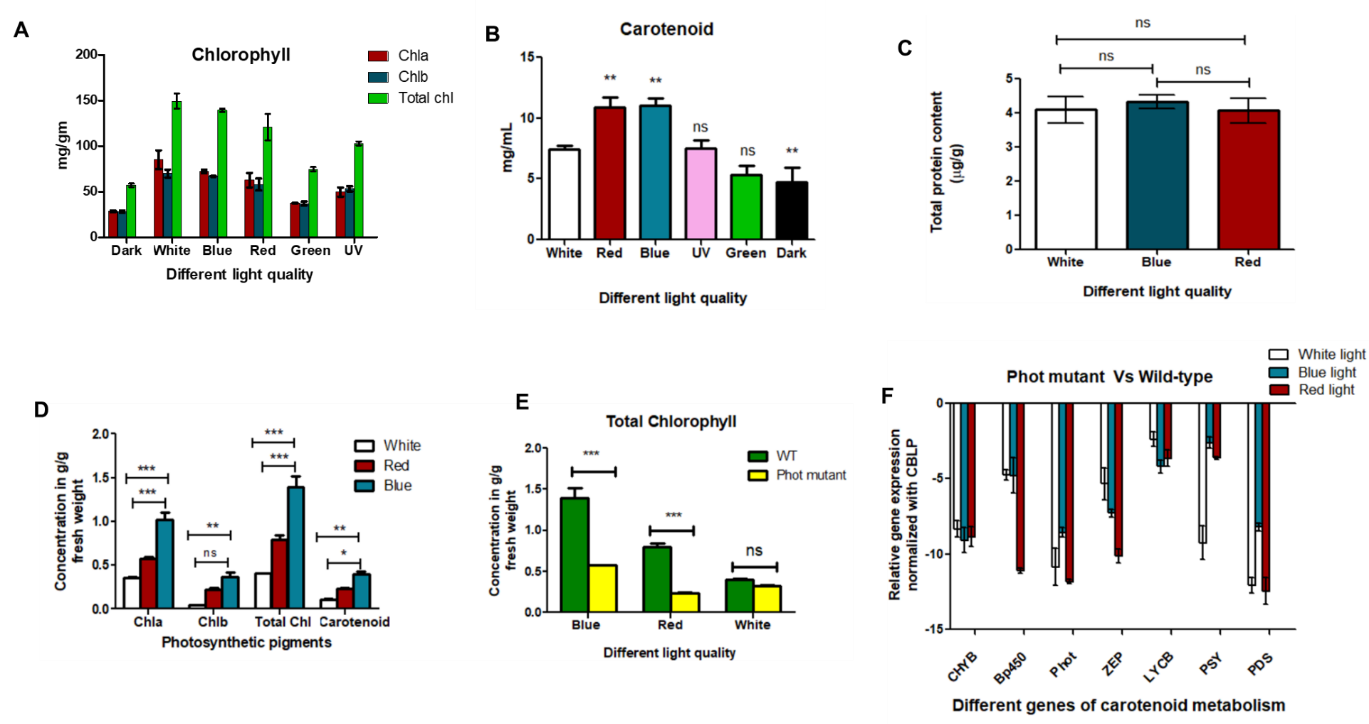
The effect of different light quality was assessed on the dark-adapted synchronized cells (12 hours light/ 12 hours dark). At the end of the dark phase the cells were exposed to different light regimes such as blue, red, green, UV, white and dark for 2 hours at approximately 4000 lux. This was followed by centrifugation at 5000 rpm for 10 min at 4℃. The pellets were used for protein isolation and metabolite extraction. The metabolite extraction was performed using 80% acetone. The total protein obtained from *Chlamydomonas* cell lysate was subjected to western blot for investigating the effect of light qualities on phototropin expression using lab raised antibody against the (Light Oxygen Voltage) LOV1 domain raised in goat. Phototropin (phot) expression was found to be increased in red and blue light with respect to white light. While the phot expression was decreased in dark and UV. Similar result was observed in photo pigment concentration. We investigated the influence of specific light regimes (red and blue light) on total protein level. It was found that the total protein level remains the same under the influence of continuous white, blue and red light in *C. reinhardtii.*

**Figure S2:** Effect of phototropin-mediated opto-modulation on photosynthetic pigments, proteins and metabolite gene expression level in *C. reinhardtii* and its phototropin mutant (CRISPR-Cas9 knockout ΔPHOT-B5 mt+). (A & B): Effect of different wavelengths on photosynthetic pigments using at 4000 lux. (C): Effect of different light quality on total protein level in *C. reinhardtii*. Values are mean ± SE (n = 4). ns represents non-significant changes. (D): Photopigment analysis of *C. reinhardtii* under red, white, and blue light at 120 µE. (E): Effect of different wavelengths of light on phototropin mutant chlorophyll contents compared with its wild-type CC-125. (F): Represents the gene expression profile of ZEP, PSY, PDS, LYCB, CHYB, BKT and bp450 in C. reinhardtii CRISPR-Cas9 phot mutant B5, after blue light and red-light treatment. Values are mean ± SE (n = 4). *P < 0.05, **P < 0.01, ***P < 0.001, ns – non-significant changes.

**Supplementary Figure 3**

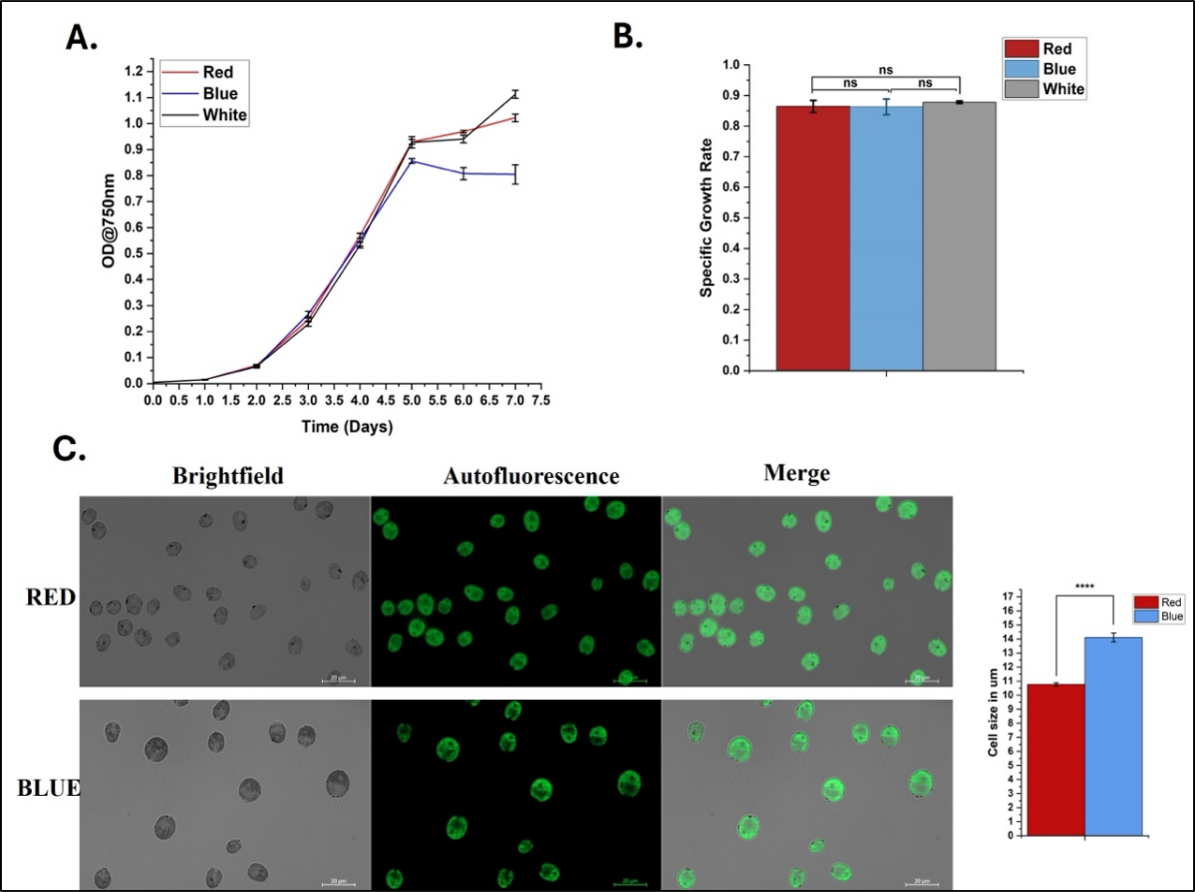
We next investigated the influence of specific light regimes (red and blue light) on growth and biomass. We observed significant effect of light on the growth and morphology of *C. reinhardtii.* Earlier studies have demonstrated that blue light delays cell division compared to red light. This might be because cultures exposed to blue light exhibited postponed DNA synthesis and cyclin-dependent kinase (CDK) activity, resulting in later cell division events Like earlier observations blue light exposure is associated with increased cell size, suggesting that cells continue to grow before division occurs, which is what we have seen here. On the contrary, red light promotes earlier cell division and may lead to a small but large number of cells. While the specific growth rates of *C. reinhardtii* cultures may appear similar under red, white, and blue light conditions, notable differences emerge during the stationary phase (Figure S3).

**Figure S3: Opto-modulation effect on growth rate and morphology of *Chlamydomonas reinhardtii*.** A) Growth curve analysis of *C. reinhardtii* under red, white, and blue light at 120 µE. (B) Specific growth rate under the three light conditions. Data represent the mean ± SE of three biological replicates. (C) Fluorescence microscopy visualization of *C. reinhardtii* cells acclimated under red (660 nm) and blue (450 nm) light. The accompanying bar graph quantifies cell size (mean ± SE) from three independent experiments, with measurements from n = 202 cells for red light (10.762 ± 0.105 µm) and n = 84 cells for blue light (14.108 ± 0.302 µm). A t-test was performed to determine the significance.

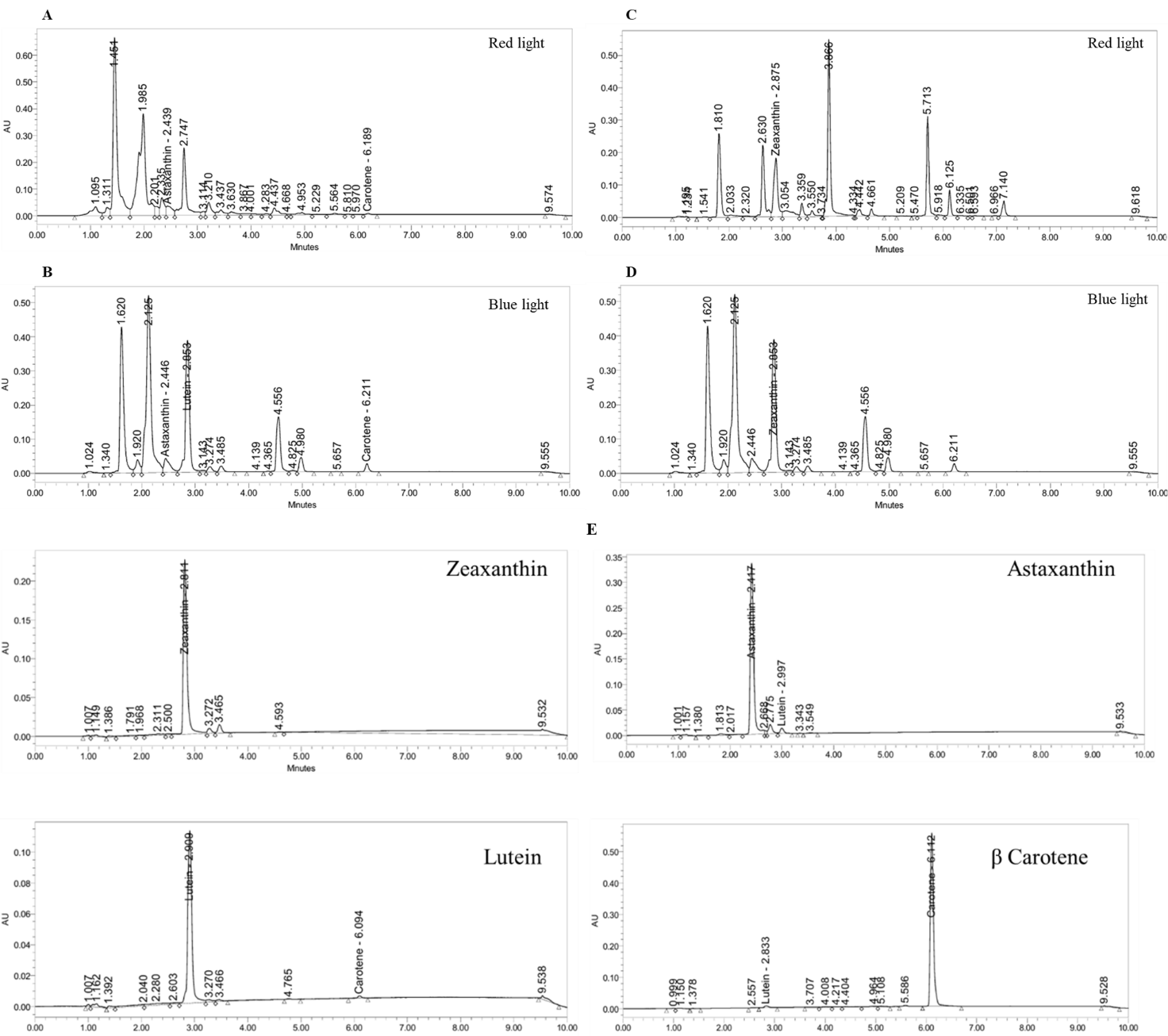
**Supplementary Figure 4**

**Figure S4: Chromatogram of the pigments and their standard (Zeaxanthin, Astaxanthin, Lutein and β Carotene) for Ultra Performance Liquid Chromatography (UPLC) used for metabolite quantification as shown in Fig. 4.** A and B: represents chromatogram for pigment astaxanthin, lutein; C and D represents chromatogram for pigment zeaxanthin. E: represents chromatogram for the standard. A total of five different concentrations of each standard was prepared. The amount was calculated based on the retention time and area of the peak obtained.

**Supplementary Figure 5**

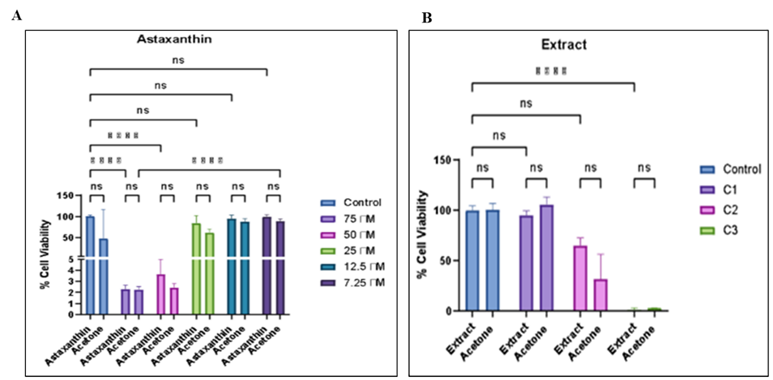
The cytotoxicity of positive astaxanthin, acetone control, and extract-treated samples on RAW 264.7 macrophage cells was evaluated using the MTT assay. Dose-dependent reduction in cell viability in different treatment groups were observed. At higher concentrations (more than 25µM), both the astaxanthin as dissolved in acetone and the acetone control exhibited reduced cell viability, indicating that the solvent contributed significantly to the observed cytotoxicity. Among the tested concentrations, lower doses (less than 25µM) of the extract maintained comparatively higher cell viability, suggesting minimal toxic effects on RAW 264.7 cells. Similarly, astaxanthin-treated cells showed acceptable viability within the selected concentration range. Concentrations that maintained≥80% cell viability were considered biocompatible and suitable for further experiments. Similarly, the cytotoxicity of the acetone extract on RAW 264.7 macrophage cells was assessed using the MTT assay. A dose-dependent decrease in cell viability was observed in different treatment. Based on these observations, only the biocompatible concentrations showing minimal reduction in viability were selected for subsequent qRT-PCR and anti-inflammatory studies to avoid interference from cytotoxic effects. Altogether, the MTT assay confirmed that the selected experimental concentrations were non-lethal and biocompatible with RAW 264.7 cells, making them appropriate for evaluating the immunomodulatory and anti-inflammatory potential of the extract. Concentrations showing ≥80% cell viability were considered biocompatible and selected for further qRT-PCR and anti-inflammatory studies.

**Figure S5:** (A) Effect of astaxanthin, extract, and acetone control on the viability of RAW 264.7 macrophage cells analysed by MTT assay. RAW 264.7 cells treated with different concentrations of astaxanthin (positive control), extract dissolved in acetone, and corresponding acetone controls were incubated for 24 h. Data is represented as mean ± SD of independent experiments. (B) Evaluation of cell viability of RAW 264.7 macrophage cells following treatment with different concentrations of algal extract and corresponding acetone controls using the MTT assay. Concentrations showing ≥80% cell viability were considered biocompatible. Values are mean ± SD of independent experiments.

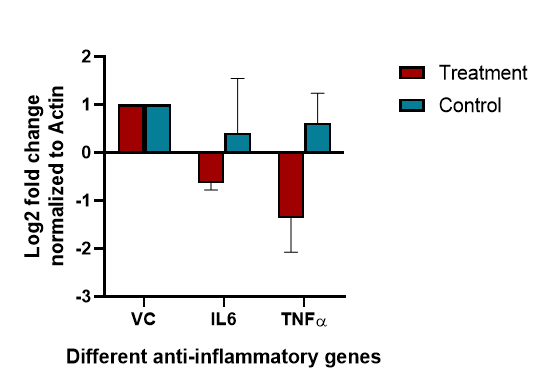
**Supplementary Figure 6**

**Figure S6:** Effect of *C. reinhardtii* astaxanthin extract under blue light on the inflammatory gene. Two biological replicates were used. Values are mean ± SE (n = 3). VC- vehicle control (acetone), IL6- Interlukein-6, TNFα- tumor necrosis factor alpha.

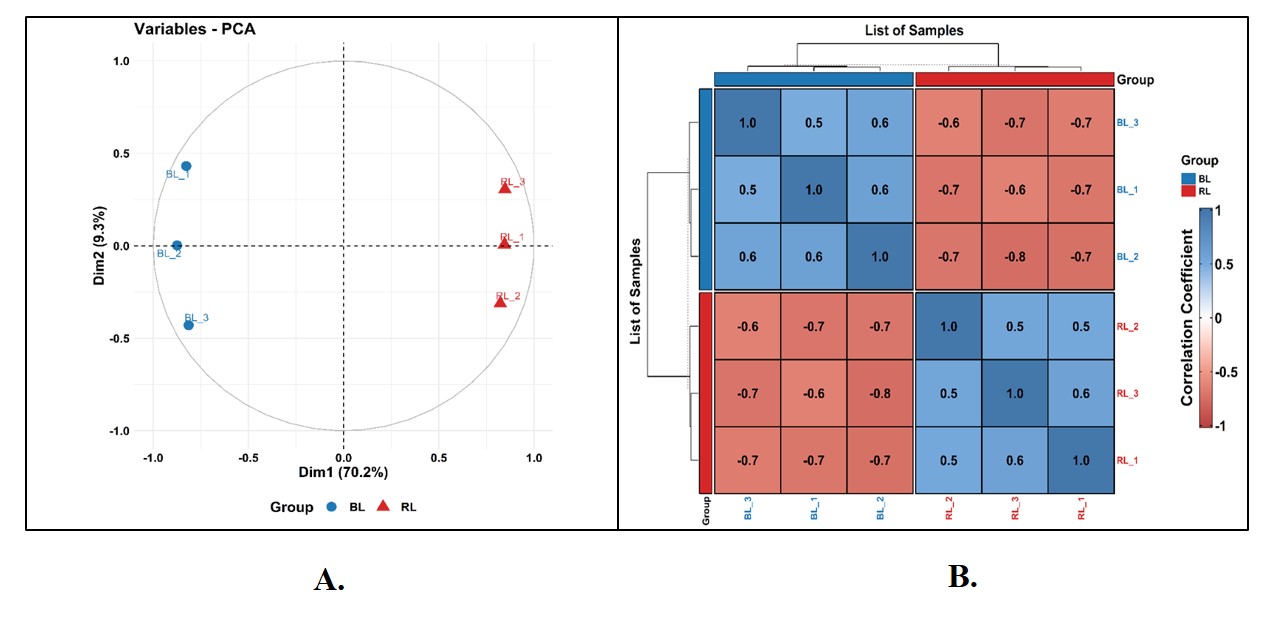
**Supplementary Figure 7**

**Figure S7: Comparative label-free quantitative proteomics analysis**

A: Principle component analysis of the BL vs RL comparative proteomics data. Principle component analysis (PCA) plot showing 3 biological replicates of BL (blue) and RL (Red). The CC-125 strain in two different light conditions cluster separately along the PC1 axis, thus, indicates differences between them at proteomic level under different light conditions. B: Represents correlation coefficient of CC-125 strain of *C. reinhardtii* under blue and red light conditions.

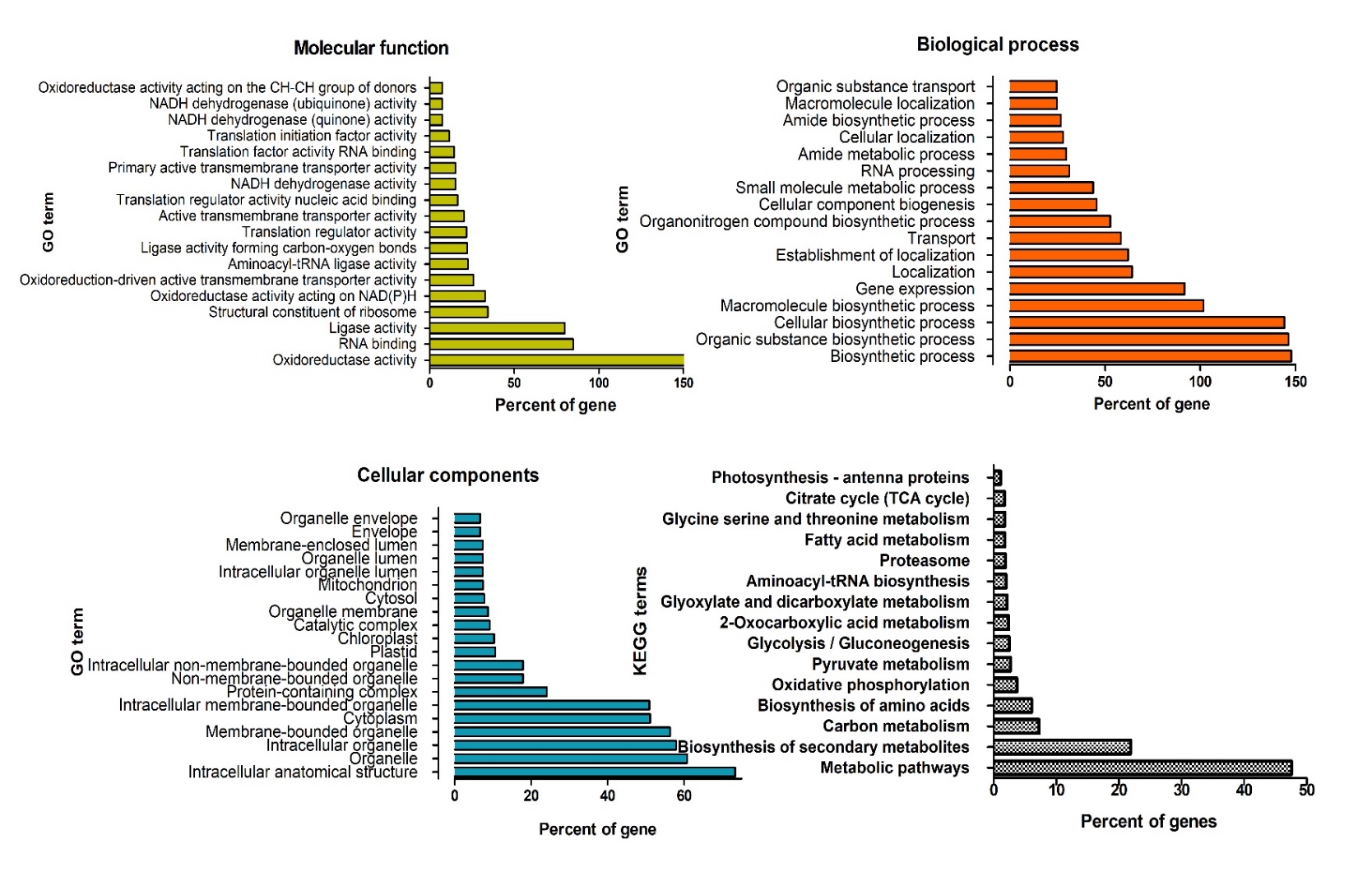
**Figure S8**

**Figure S8: The figure represents top 20 GO terms based on cellular components, molecular function, biological function and percent of genes linked to each GO term in *C. reinhardtii* in response to blue light.**

The gene percent was calculated by using term size and intersection size of the DEPs based on their rank.

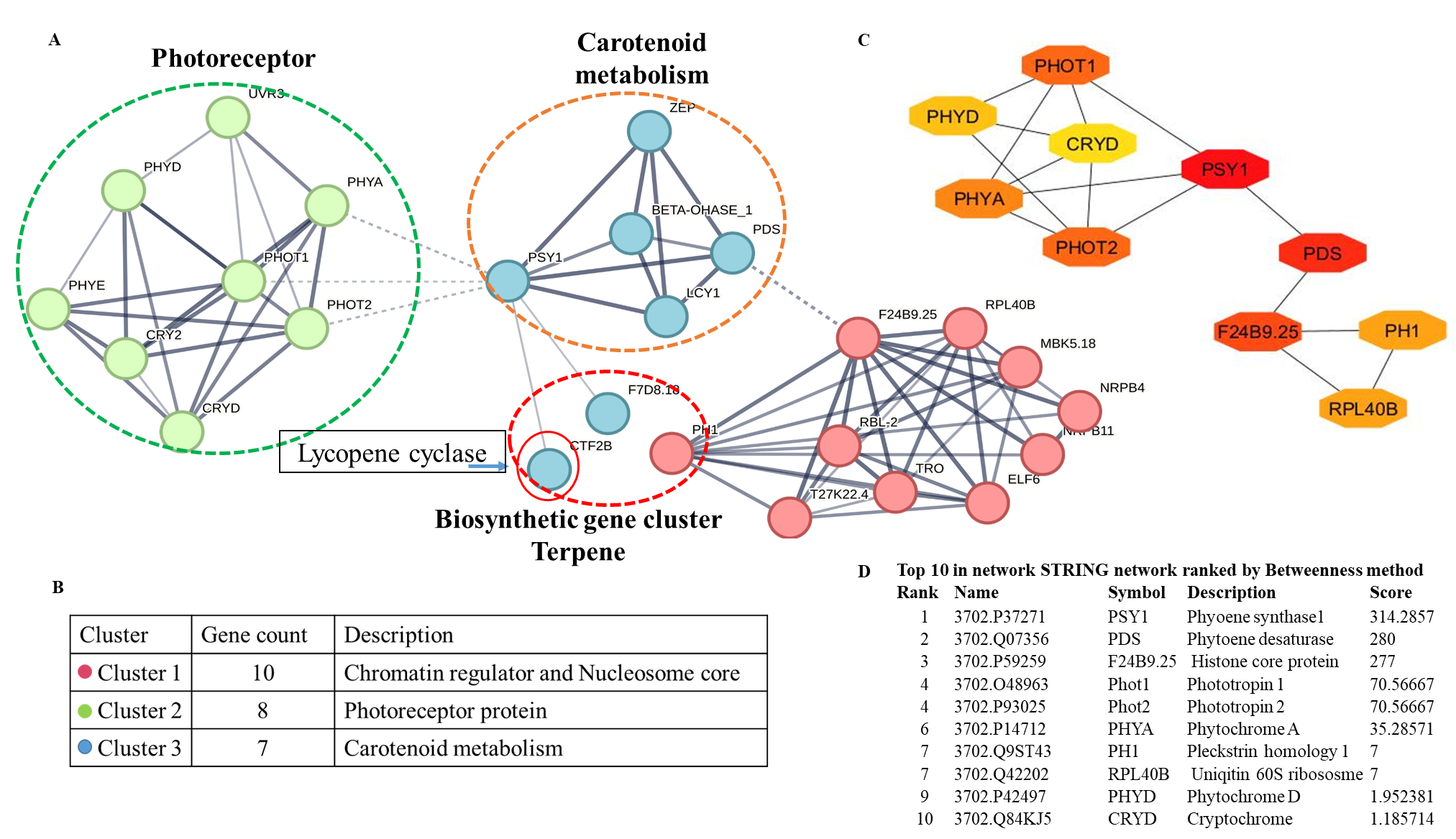
**Supplementary Figure 9**

**Figure S9: Bio-curation of opto-modulation of BGCs for targeted metabolite production in *Arabidopsis thaliana***

**A:** It shows the bio-curated crosstalk of BGCs cluster and photoreceptor via predicted curated PPI networking in *A. thaliana*. Here, lycopene cyclase (BGC core domain component showed in red circle) showed direct interaction with phytoene synthase 1 (a key enzyme of carotenoid metabolism). **B:** The k-means clustering method was applied to cluster the PPI. The secondary interaction of photoreceptors was observed with lycopene cyclase protein biosynthesis BGCs component. **C:**  It represents the network analysed based on top 10 betweenness methods and shortest path methods. Colour range (red to yellowish) indicates scores ranked by the betweenness method with their interacting partners. **D:** The rank score is represented in lower panel.

**Supplementary Figure 10**

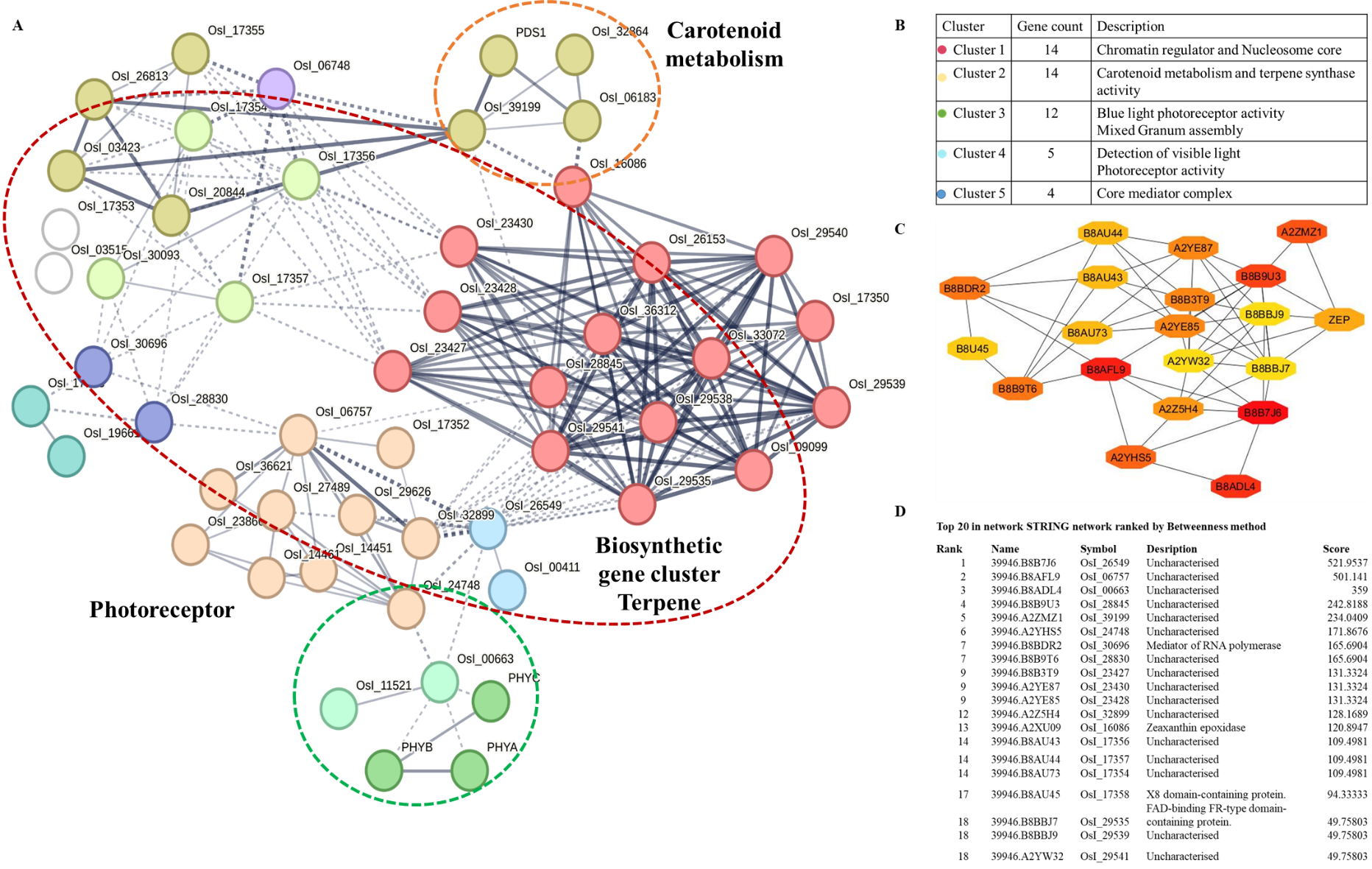

**Figure S10: Bio-curation of opto-modulation of BGCs for targeted metabolite production in *Oryza sativa (indica)***

**A:** It shows the bio-curated crosstalk of BGCs cluster and photoreceptor via predicted curated PPI networking in *A. thaliana*. Here, the secondary interaction of photoreceptors was observed with terpene biosynthesis BGCs component. **B:** The k-means clustering method was applied to cluster the PPI. **C:**  It represents the network analysed based on top 10 betweenness methods and shortest path methods. Colour range (red to yellowish) indicates scores ranked by the betweenness method with their interacting partners. **D:** The rank score is represented in lower panel.

**Raw Blot Images**

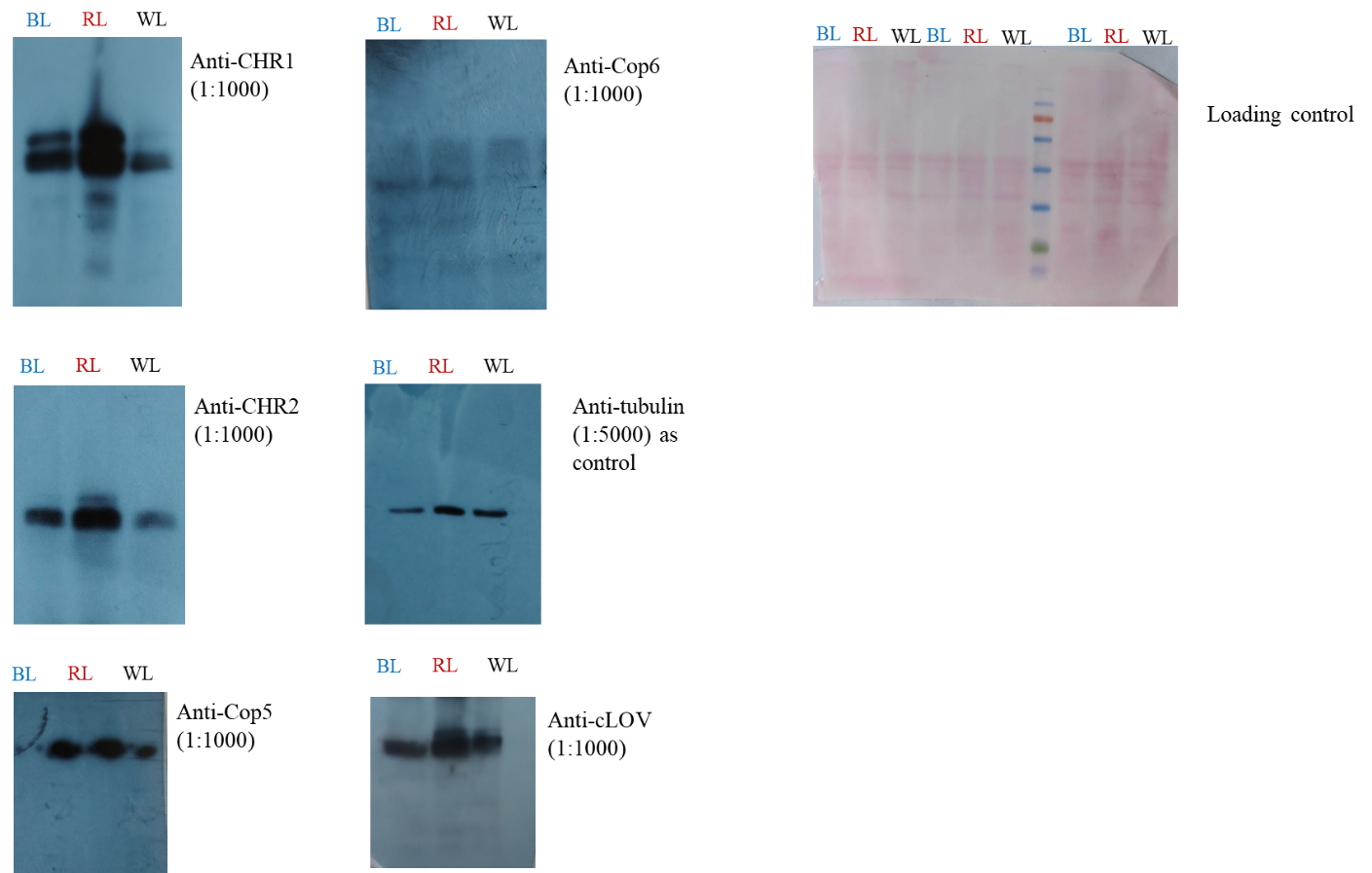
